# ESX-5 Deletions in *Mycobacterium tuberculosis* Alter Macrophage Cytokine Signaling and Bacterial Heavy Metal Response

**DOI:** 10.1101/2025.09.30.679525

**Authors:** Austin M. Haynes, Sara B. Cohen, Aparajita Pesaladinne, David R. Sherman, Kevin B. Urdahl, Jeffery S. Cox, Thomas R. Hawn

**Author notes:** Corresponding Authors: Thomas R. Hawn & Austin M. Haynes.

## Abstract

The ESX-5 secretion system is critical for *Mycobacterium tuberculosis* (Mtb) viability and putatively linked to pathogenesis, but a functional understanding of how it interacts with the host is unknown. ESX-5 is encoded at a single genomic locus with small, paralogous, secreted targets (ESX-5a, 5b, 5c) spaced throughout the genome. To examine host-pathogen interactions of these putative virulence clusters, we made mutant strains lacking these loci and infected primary human macrophages. Surprisingly, all deletion mutants independently reduced cytokine secretion during infection, specific to certain analytes. This defect depended on viable bacteria and was mediated by a post-transcriptional mechanism. In bacterial transcriptomic analyses, each mutant downregulated heavy metal response genes compared to wild type bacteria. Treatment of Mtb with Cu or Cd led to increased ESX-5a and ESX-5c expression, concurrent with increased TNF and IL-6 secretion in macrophages compared to untreated bacilli, indicating a link between ESX-5 expression and cellular cytokine levels.

## Introduction

Humans and *Mycobacterium tuberculosis* (Mtb) have co-evolved for thousands of years shaping host and pathogen genomes, facilitating survival advantages, and causing extensive disease burden (1–6). Via specially adapted virulence systems Mtb can block phagolysosomal maturation, inhibit phagosome acidification, modulate autophagy, and induce macrophage necrosis, among other processes (7). The molecular details of how Mtb carries out many of its virulence functions are not fully described, particularly during early infection when the bacillus is phagocytosed by macrophages (7–9). Elucidating a more complete understanding of the interplay between bacterial effectors and human macrophages could lead to the development of novel therapeutic targets. Mtb contains several bacterial secretion systems that enable translocation of these effector molecules to host interfaces with the potential to modulate antimicrobial pathways (10–13). These systems include the general secretion (SEC), twin arginine translocation (TAT), and type seven secretion systems (T7SS), which are also called ESX (14–17). Mtb encodes five ESX systems, denoted ESX-1 through ESX-5, which secrete roughly 5-10% of Mtb encoded proteins (16,18–21). Some ESX systems have predicted or known virulence function, such as CFP-10 and ESAT-6 via ESX-1 (22–24). Others, like ESX-5, are less well defined.

ESX-5 secretes many proteins, including members of the Proline-Glutamic Acid (PE), Proline-Proline-Glutamic Acid (PPE), Esx, PE-Polymorphic GC-Rich Repetitive Sequence (PE_PGRS), and PPE- Major Polymorphic Tandem Repeat (PPE_MPTR) families (25–27). Together, these families comprise >100 proteins and are extensively expanded in the Mtb genome (28–30). Although functional characterization is lacking for most of these effectors, many are thought to be involved in immune subversion, nutrient acquisition, membrane structure, and virulence (29,31–36). Furthermore, recent clinical and genetic studies have specifically implicated paralogous ESX-5 secreted clusters in virulence, transmissibility, and immune recognition (37–40). These paralogs are each comprised of a four gene cluster and named ESX-5a, ESX-5b, and ESX-5c respectively. While previous studies suggest an important role for ESX-5 in TB pathogenesis, the molecular and cellular details remain poorly defined, particularly with regards to interactions with macrophages.

To begin addressing these gaps in knowledge, we created targeted deletions of ESX- 5a, 5b, and 5c to investigate the role of these paralogs during human macrophage infections. We queried differential responses to the knockouts in vitro and in axenic culture to develop a model of the role ESX-5 plays in modulation of innate immune responses to Mtb and bacterial physiology.

## Materials and Methods

### Ethics and Biosafety Precautions

Primary human monocyte derived macrophages (MDMs) from healthy human donors in the Seattle area were from Bloodworks Northwest and are deidentified prior to receipt. No IRB is required to perform studies with these cells. All experiments performed with live, virulent, Mtb were performed in an accredited BSL-3 at the University of Washington School of Medicine campus utilizing validated containment and inactivation protocols to ensure biological safety.

### Bacterial Strains and Cultivation

H37Rv (Sherman) used in our laboratory was a gift from Dr. David Sherman. H37Rv pNIT was received from Drs. Christoph Grundner and Andrew Frando from Seattle Children’s Research Institute (SCRI). All knockout strains (ΔESX-5a, ΔESX-5b, and ΔESX-5c) are descended from the H37Rv pNIT strain of MTB. Complementation strains are decedents of their respective knockout parent strain. All CRISPRi strains used in this study (EccC5i) are descended from H37Rv (Sherman) strain. To culture these MTB strains, a frozen stock was thawed to room temperature and added to 4mL of 7H9-GAT comprised of 7H9 base along with 0.2% Glycerol (**G**), 10% ADC supplement (**A**) and 0.05% Tween 80 (**T**). Appropriate selection was added if required. Strains were cultured shaking and vented until OD reached ∼0.8-1.0, then back diluted to OD 0.1, and outgrown to mid-logarithmic phase growth before using in macrophage infections.

Strain List:

H37Rv (Sherman)

H37Rv (Sherman) EccC5i (EccC5 CRISPRi Knockdown)

H37Rv pNIT

H37Rv pNIT ΔESX-5a (ΔESX-5a) (deletion of PE8/PPE15/EsxI/J)

H37Rv pNIT ΔESX-5b (ΔESX-5b) (deletion of PE13/PPE18/EsxK/L)

H37Rv pNIT ΔESX-5c (ΔESX-5c) (deletion of PE32/PPE65/EsxV/W)

H37Rv pNIT mCherry

H37Rv pNIT ΔESX-5a (ΔESX-5a) mCherry

H37Rv pNIT ΔESX-5b (ΔESX-5b) mCherry

H37Rv pNIT ΔESX-5c (ΔESX-5c) mCherry

ΔESX-5a pTEC15 ZeoR::ESX-5a Comp.

ΔESX-5a pTEC15 ZeoR::EsxI/J Comp.

ΔESX-5a pTEC15 ZeoR:: PE8/PPE15 Comp.

ΔESX-5b pTEC15 ZeoR::ESX-5b Comp.

ΔESX-5b pTEC15 ZeoR::EsxK/L Comp.

ΔESX-5b pTEC15 ZeoR::PE13/PPE18 Comp.

ΔESX-5c pTEC15 ZeoR::ESX-5c Comp.

ΔESX-5c pTEC15 ZeoR::EsxV/W Comp.

ΔESX-5c pTEC15 ZeoR::EsxV/W T2A Comp.

ΔESX-5c pTEC15 ZeoR::PE32/PPE65 Comp.

### pNIT Allelic Exchange

Recombineering was performed using RecET mediated homology directed exchange as described previously(41). Briefly, we PCR amplified 500 base pair homology arms flanking our Mtb genome regions of interest, and generated a PCR product of a hygromycin resistance marker under control of the mycobacterial optimized promoter (MOP). Concurrently, we linearized our destination plasmid pUC19 within the MCS, performed an InFusion reaction (Takara Bio, San Jose CA) as described by the manufacturer, transformed Stellar Competent Cells (Takara Bio, San Jose CA), isolated colonies, purified plasmids, and validated using whole plasmid sequencing. From this stock construct, our linear exchange fragment was amplified using PCR flanking the 5’ and 3’ ends. After generating linear exchange template, we then used standard MTB electroporation protocols to introduce the exchange construct. Initially we induced the expression of the gp130 and 131 genes within Mtb H37Rv containing the pNIT plasmid, using Isovaleronitrile (sigma Aldrich, Saint Louis MO). We then made these recombinogenic bacteria electrocompetent via pelleting and successive washing using a chilled 10% glycerol solution. The final pellet was resuspended in 3mL (10% initial culture volume) 10% glycerol. 400uL of competent cells were added to a 2mm gap cuvette (BioRad, Hercules CA) with ∼1 microgram of exchange template then electroporated using an exponential pulse of 2.5 kV, 1,000 Ω, and 25 μF. Bacteria were immediately recovered into 5mL of 7H9-GAT overnight, then plated overnight on selective 7H10 agar for outgrowth. Outgrowth proceeded for a maximum of 5 weeks. Colonies were selected and screened using molecular and phenotypic methods.

### CRISPRi Strain Generation

CRISPRi plasmid cloning for target guides was performed as described previously(42). We identified candidate guide sequences in the *eccC5* gene using an in-silico approach to determine strong PAM sites. After electroporating the CRISPRi plasmid into Mtb and determining successful integration via phenotypic testing, we induce the expression of the CRISPRi machinery using anhydrotetracycline (ATc). Once induction reached maximum levels, we infected MDMs as described.

### Complementation vector cloning

Complementation cloning was performed using the pTEC15 plasmid (Gift of Dr. Rafael Hernandez). Complementation constructs were generated using a homology directed cloning method. First, pTEC15 was linearized using the EcoRV restriction enzyme (New England Bio Labs, MA), destroying the mWasabi reporter marker. Insertion DNA was amplified using sequence specific primers containing 15 base pair homology arms for the restriction site used. Insertion DNA and linear vector were then ligated using homology directed recombination with the InFusion system (Takara Bio, San Jose CA). These plasmids were then validated using whole plasmid sequencing.

### Macrophage Generation and Column Isolation

Human peripheral blood mononuclear cells (PBMC) were obtained from Bloodworks Northwest. We obtain leukocyte reduction system (LRS) cones that remove leukocytes during blood donation and extract cells from this cone via standard Ficoll gradient isolation described previously(43). Isolated PBMC are stored in liquid nitrogen at 50E6 cells per vial until use. To generate macrophages, a PBMC stock was drawn from liquid nitrogen storage and thawed in a 37C water bath. One milliliter of RPMI-10 was then added to the stock to bring up temperature while concurrently diluting DMSO of the freeze media. This mixture was then added to 5mL of RPMI-10 and pelleted at 1250g for 5 min. The cells were washed x2 using RPMI-10 before final resuspension in RPMI- 10 in a non-tissue culture treated petri dish. Cells were incubated for 5-7 days at 37C and 5% CO2. At the end of incubation, macrophages were harvested by manual scraping and isolation via CD14+ affinity isolation using the MACS column separator according to manufacturer’s protocol (Miltenyi Biotec, Germany). Cells are then plated at varying concentrations and rested overnight at 37C, 5% CO2 before infection the following day.

### Macrophage Infections

At the time of infection, bacterial culture was harvested via centrifugation (4000rpm, 5- 10 min) and washed x2 with Sautons medium. The cultures were then resuspended in 2.5mL of Sautons medium and enumerated using OD600 absorbance and a standard conversion factor of 5E8 CFU/mL at OD600 of 1.0. After enumeration, necessary volumes of the bacterial suspensions were diluted into RPMI-10 to achieve the necessary MOI for the experiment. This media was then used to exchange the media on the macrophages. To bring bacteria into contact with cells rapidly, a spin at 500g for 5 minutes was performed.

### Alamar Blue Assay

Alamar Blue reagent was added at 10% v/v to each well of a plate minus the blank wells as a media control. Cells were infected as described and fluorescence read (570nm excitation, 585nm emission) at 0 hours post infection (hpi) to establish baseline fluorescence levels. The reaction was allowed to proceed overnight with the next florescence measurement being collected at 18hpi. Uninfected wells were set as the baseline to determine the lower range of redox within this cell type, infected conditions were compared to this group to evaluate viability (higher vs. lower redox). 48hpi read was performed the following day to determine deterioration rates of fluorescent signal at which point assay was terminated.

### Enzyme Linked Immunosorbent Assay (ELISA)

DuoSet (R&D Systems Minneapolis MN) enzyme linked immunosorbent assay (ELISA) was performed for all ELISA assays per manufacturer’s instructions.

### LEGENDplex Cytokine Analysis

Staining was performed according to the manufactures protocol with a custom analyte panel (LEGENDplex, biolegend.com). Supernatants were isolated from infected cells and added to a bead mixture containing capture beads for our target analytes. Flow analysis identified analyte bead populations based on color and bead size. We analyzed using BioLegend software where standard curves were calculated for respective analytes.

### Interferon Beta Reporter Assay

Supernatants from infected macrophages (containing IFNβ) were used to treat a transgenic Huh7 cell line containing an MX1 linked gaussia luciferase gene with an ISGF3-linked IFNAR (gift from Dr. Ram Savan)(44). Within the Huh7 cell line, binding of IFNAR to IFNβ leads to the downstream transduction of ISGF3 to the MX1 promoter element. MX1 activation leads to expression and secretion of a codon optimized gaussia luciferase (GLuc). After 24 hours of IFNβ stimulation on this transgenic cell line, supernatants were collected, mixed 1:1 with a coelenterazine substrate (ThermoFisher, Waltham MA) and used to measure luminescent signal.

### Acidic Phagosome Staining

LysoTracker Deep Red (L12492) (ThermoFisher, Waltham MA) was used to selectively label acidified compartments within cells. Macrophages were column isolated, plated, rested overnight, and infected with H37Rv or knockout strains at an MOI of 1.0. for four hours before adding LysoTracker at a final concentration of 100nM. LysoTracker treatment was run for two hours followed by a media exchange with RPMI-10. Treated cells were then measured via Mean Fluorescence Intensity (MFI) at 647nm excitation and 668nm emission using a BioTek Cytation 5 (Agilent, Santa Clara CA).

### Macrophage Fractionation

Cells were fractionated to collect paired secreted and cellular fractions from the same sample and query cellular cytokine concentrations. Cells were infected as described. At the time of harvest, cellular supernatants were collected and filtered in a 0.22-micron PVDF low protein binding filter plate. For the residual monolayer, it was washed x2 with 1x HBSS to remove any residual supernatant. Following wash, the monolayer was lysed with 1% Triton x-100 in water solution (containing protease and phosphatase inhibitors (Roche c0mplete tabs)) at room temperature for 10-20 minutes. The lysate was then agitated using a multichannel pipette to ensure complete lysis and visualized via light microscopy before filtration in a 0.22-micron low protein binding filter plate.

### MG132 & UPR/ISR Inhibitor Treatment

MG132 proteasome inhibitor (Millipore-Sigma, Burlington MA) was diluted to a final concentration of 10µM in RPMI-10. Media was then exchanged onto cells at the start of Mtb infection and remained for 24 hours. Concentration determined from previous literature(45). UPR and ISR inhibitors (Selleck Chemicals (Houston, TX)) were tested against macrophages on a dose curve to determine maximal dosing before toxicity occurred. For each infection, each inhibitor was then diluted into RPMI-10 at their respective experimental concentrations, 4µ8c at 15µM, Ceapin A7 at 10µM, GSK at 1µM, and ISRIB at 250nM. Inhibitors were added to cells ∼2 hours before infection. Per respective infections, bacteria were then spiked in to achieve MOI of 5. After 24 hours, supernatants were collected and analyzed via ELISA as described.

### CFU Assay

MDMs were infected at an MOI of 1.0 and spun at 500g for 5 minutes. After 72 hours of infection, media was removed from the monolayers, and the cells were lysed with 1% Triton X-100 solution in water. The lysate was then serially diluted 10-fold in 7H9-GAT containing Tween to disperse bacterial clumps for a final dilution range of 10^-1^ to 10^-6^ for all samples. 100uL of the 10^-3^ through 10^-5^ was plated on 7H10 agar plates and dried at room temperature. Plates were outgrown for ∼2 weeks and colonies enumerated.

### Macrophage and Mtb gDNA Isolations, RNA Isolations, and Gene Expression Quantitation

Genomic DNA was extracted from Mtb via a crude lysate method. Cultures were grown to an optical density of 1.0, pelleted at 14,000 rpm for 10 minutes, resuspended in water, and boiled at 95C for 30 minutes to inactivate and lyse the bacteria. After 30 minutes, the lysates were pelleted again to remove insoluble debris and filtered through a 0.22-micron Polyether Sulfone (PES) filter unit for additional purification. To quantify gene expression of target genes in macrophages and Mtb we performed a probe-based gene expression assay approach. For macrophages, RNA was isolated using a Trizol isolation technique. We resuspended cell monolayers in Trizol and isolated RNA using the manufactures protocol paired with an RNeasy Extraction kit (Qiagen, Hilden Germany). For Mtb, we resuspended bacteria in Trizol (Invitrogen, Waltham MA), transferred to Lysing Matrix B tubes (MP Biomedicals, Santa Ana CA), homogenized the sample using a 6500rpm, 30 second protocol on a beadruptor (Omni International, Kennesaw GA) followed by a 30 second incubation on ice. This was repeated x3 to ensure complete lysis of the bacteria and RNA liberation. We then performed RNA isolation the same as described above. Mtb or macrophage RNA was reverse transcribed with a cDNA generation protocol using a high-capacity cDNA kit (Life Technologies, Waltham MA). Resulting cDNA was used in a probe-based, junction spanning (for Macrophages), gene expression assay reaction to quantify transcript levels (IDT, Coralville IA). For macrophage studies, target-specific probe sets were multiplexed with an endogenous control probe targeting human GAPDH. For Mtb studies, the control was sigma factor A (SigA). Relative quantitation was performed using the ΔΔCt method.

### Mtb RNA Sequencing

Total RNA was extracted from Trizol samples and ribosomal RNA (rRNA) was depleted using a previously described protocol(46). In brief, biotinylated DNA probes were generated with complementarity to the ribosomal RNA sequences for Mtb. These probes target 46 different rRNA species, covering 23s, 16s, and 5s rRNA sequences (**Supplemental Table 1**). Probes were mixed with total RNA and hybridized using a decreasing temperature gradient approach. Streptavidin magnetic beads were added to the hybridized mixture and incubated to bind the biotin moiety on probes, then isolated using a magnetic separator. The supernatant containing rRNA depleted RNA was then cleaned further using AMPure XP beads (Beckman Coulter, Brea CA). Bead-bound RNA was washed to remove impurities prior to elution in TE buffer. Purified RNA was then prepared for paired end 150 base pair short read sequencing (PE150) using the NEBNext Ultra II RNA Library Prep Kit for Illumina (New England Biolabs, Ipswich, MA). Libraries were submitted to Northwest Genomics Center (NWGC) sequencing core for library QC and sequencing. Quality was assessed using a High Sensitivity DNA Assay for the Bioanalyzer 2100 (Agilent, Santa Clara CA). All libraries were then multiplexed and sequenced using an Element Aviti (Element Biosciences) 300 Cycle Low Output chip totaling 2.5E8 reads.

### Bioinformatics Analysis

Raw FASTQ data from the sequencing core was initially inspected using the FASTQC program to visualize read length, read quality, among other quality metrics prior to further analysis. After initial quality control, the raw sequences were run through our labs SEAsnake pipeline for RNA sequencing data processing (47). Adapter sequences were trimmed, low-quality reads removed using the “AdapterRemoval” program and sequences were aligned to reference using the STAR program. Alignment filtering and output quality were assessed using samtools (“flagstat”) and gene counts were generated with the “Subread” package. Differentially expressed genes were calculated across our samples, averaging experimental replicates for statistical analysis, using the DEseq2 package(48). Differentially expressed genes run through an inclusion threshold using FDR correction and log2 fold change relative to wildtype baseline (FDR≤0.1, Log2FC≤1). Output gene lists were intersected to identify unique and overlapping gene hits across our strains. These lists were then used in a string analysis (StringDB) to identify clustering with concurrent hypergeometric enrichment analysis (PantherDB) on Gene Ontology (GO) Biological Processes gene set database.

### Culture of Mtb in Metal-containing Medium

Cultures were grown as described then back diluted into four separate rich medium cultures tubes containing heavy metal (Cu, Cd, Zn, or Fe). For growth curves, Cu and Cd cultures were treated with 100µM of the respective metal while Zn and Fe cultures were treated with 500µM. Expression analysis was performed on cultures treated for 18 hours with high dose Cu or Cd (1000µM) before RNA harvest. For MDM infections, metal pretreatment of these bacteria was performed as outlined above with H37Rv grown in low dose Cu or Cd containing 7H9-GAT (100µM of respective metals) for 5 days and then subject to standard washing and infection protocol. After 24 hours, cellular supernatants were isolated and filtered. These supernatants were then evaluated using ELISA.

## Results

### MDM infection with Mtb ESX-5 knockout strains results in reduced proinflammatory cytokine secretion

To assess the role of ESX-5 globally in Mtb-infected macrophages, we used CRISPR interference to target EccC5, an essential ATPase that regulates secretion(49).

Knockdown of this gene was associated with decreased growth in primary human MDMs, but not in log phase broth culture (**Supplemental Figure 1**). We next deleted three ESX-5 paralogs (**Supplemental Figure 2A-B)** and examined the impact of these mutants (ΔESX-5a, ΔESX-5b, and ΔESX-5c) on MDM responses during early infection by assessing Mtb replication in vitro and in axenic culture, acidified phagosome content, and proinflammatory cytokine output. We observed no difference in bacterial growth in broth culture or in MDMs for all strains tested (**Figure 1A-B**). The level of acidified phagosome content was slightly lower in ΔESX-5c, but not ΔESX-5a or ΔESX-5b infected macrophages, relative to H37Rv (**Figure 1C**). TNF, IL-6, and IL-1β were all significantly reduced in the secreted fraction of knockout Mtb-infected cells compared to wild type (**Figure 1D**, MOI=5, 24-hour time point). We assessed the viability of these macrophages using an Alamar Blue assay and determined there was no difference at 24 hours post infection (hpi), indicating the differences in cytokine secretion is not due to increased cell death (**Supplemental Figure 2C**). For all three mutants, reintroduction of the cognate genes complemented the phenotype and restored cytokine secretion at or above wild type levels (**Supplemental Figure 3**). Furthermore, paraformaldehyde fixation of the knockout strains also restored cytokine secretion (**Figure 1E**). Taken together, these data suggest that ESX-5a, 5b, and 5c specifically regulate cytokine output in MDMs via a process requiring bacterial viability.

**Figure 1.**
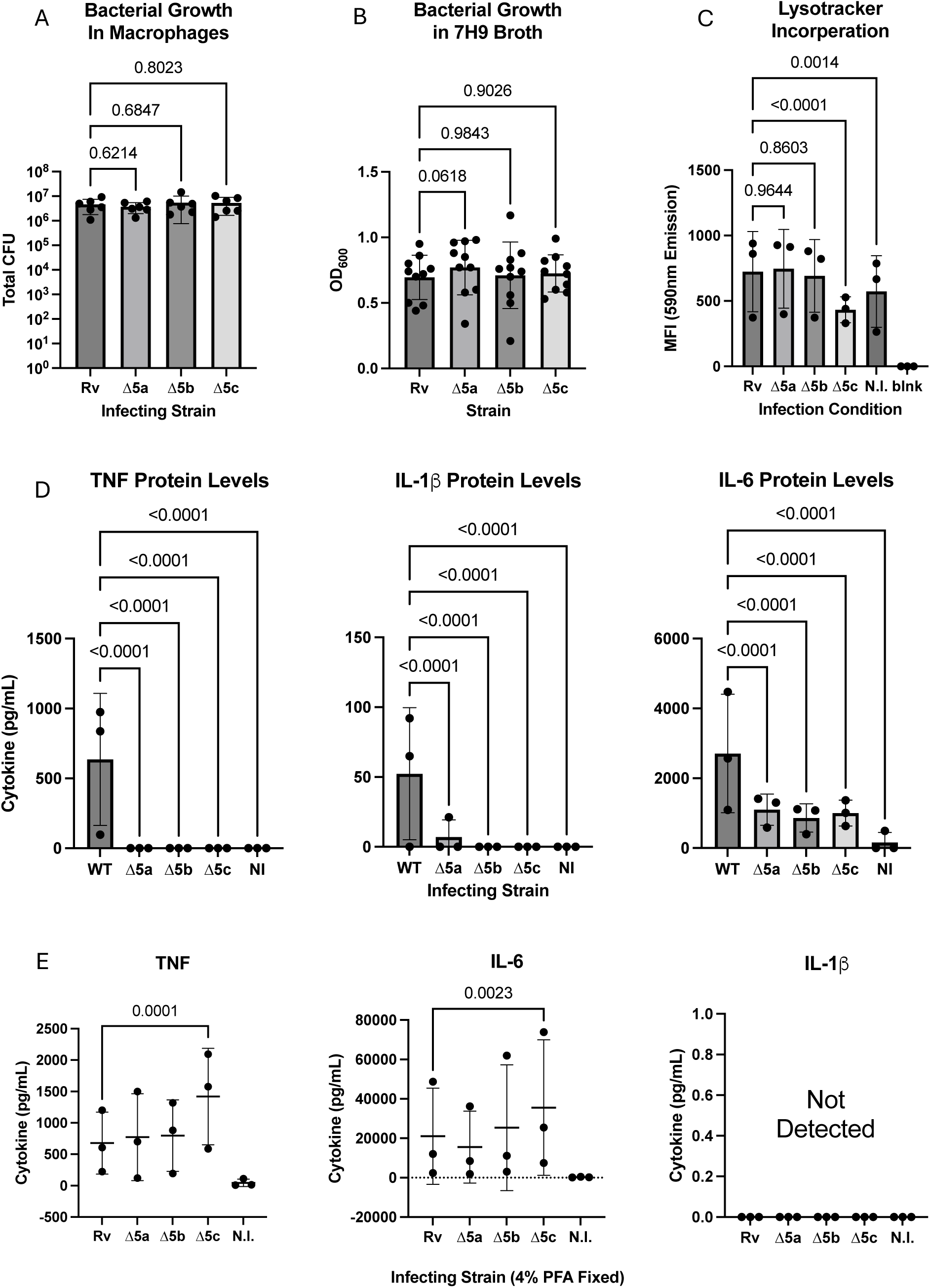
ESX-5 paralog deletion mutants induce differential inflammatory profiles in infected monocyte-derived macrophages. *1A. ESX-5 paralog deletions do not impact bacterial fitness in macrophages*. 72hr replication levels via CFU (y-axis Log10 scale) recovered from N=6 infected donors. *1B. ESX-5 deletion growth is not attenuated in broth culture*. Bar chart displays broth replication levels (y-axis OD600) for all knockout and wildtype strains (x-axis) for N=10 experiments. *1C. ESX-5 deletions alter phagolysosomal acidification.* Bar chart displays acidic organelle staining in macrophages 6 hours post infection (6hpi) for N=3 biological donors (technical triplicate). Strain used on the x-axis (N.I. = uninfected control, No LT = no lysotracker control) and mean fluorescence intensity (MFI) on the y-axis. *1D. ESX-5 deletions induce altered cytokine profile in MDMs*. Supernatant levels of TNF, IL-6, IL-1β (y-axis pg/ml) after Mtb infection of MDMs (x-axis strain or uninfected (NI)) (N=3 donors run in technical triplicate).*1E. Bacterial inactivation restores cytokine production.* Supernatant levels of TNF, IL-6, IL-1β (y-axis pg/ml) from MDMs (N=3 donors in technical triplicate) infected with 4% PFA inactivated strains or uninfected (N.I.) (x-axis). *1A,1C-E.* Statistics calculated using two-way ANOVA with a post-hoc T-test and Dunnett correction for multiple comparisons (95% confidence interval (CI), p≤0.05). *1B.* Statistics calculated using one-way ANOVA with a post-hoc T-test and Dunnett correction for multiple comparisons (95% CI, p≤0.05). Error bars show standard deviation of the mean.

### ESX-5 deletions do not alter macrophage phagocytosis, cytokine mRNA transcription, cytokine secretion, or degradation

To test potential cytokine reduction mechanisms, we first examined whether phagocytosis levels were different between our mutant and wildtype strains. We observed no difference in the number of bacteria phagocytosed at 4hpi (**Figure 2A**). Furthermore, TNF, IL-6, and IL-1β mRNA levels were not impacted by these deletion mutants at 6hpi, indicating that they retained the ability to activate innate immune signaling pathways (**Figure 2B**, MOI=5). For a subset of strains tested, we also observed no difference in RNA levels at 24hpi (**Supplemental Figure 4**). To test whether there was a secretion defect, we collected cellular and secreted fractions from infected cells and measured cytokine protein levels. Cellular lysate fractions did not contain elevated levels of cytokine in the ΔESX-5a, ΔESX-5b, or ΔESX-5c conditions. Although we could not detect intracellular TNF, IL-6 and IL-1β protein levels were markedly decreased compared to infections with wild type Mtb. This result indicated that ESX-5 inhibits host translation, regulates transcript post-transcriptionally, or increases cytokine degradation (**Figure 2C**). Finally, to test whether these differences were due to proteasomal degradation of intracellular cytokines, we added the MG132 proteosome inhibitor to the cells at the time of infection. At 24hpi, we observed no difference between wild type and knockout infected cells comparing MG132 treated to untreated controls (**Figure 2D**). Together, these data suggest that ESX-5 deletions may impact transcriptional or translational control of cytokine synthesis within macrophages.

**Figure 2.**
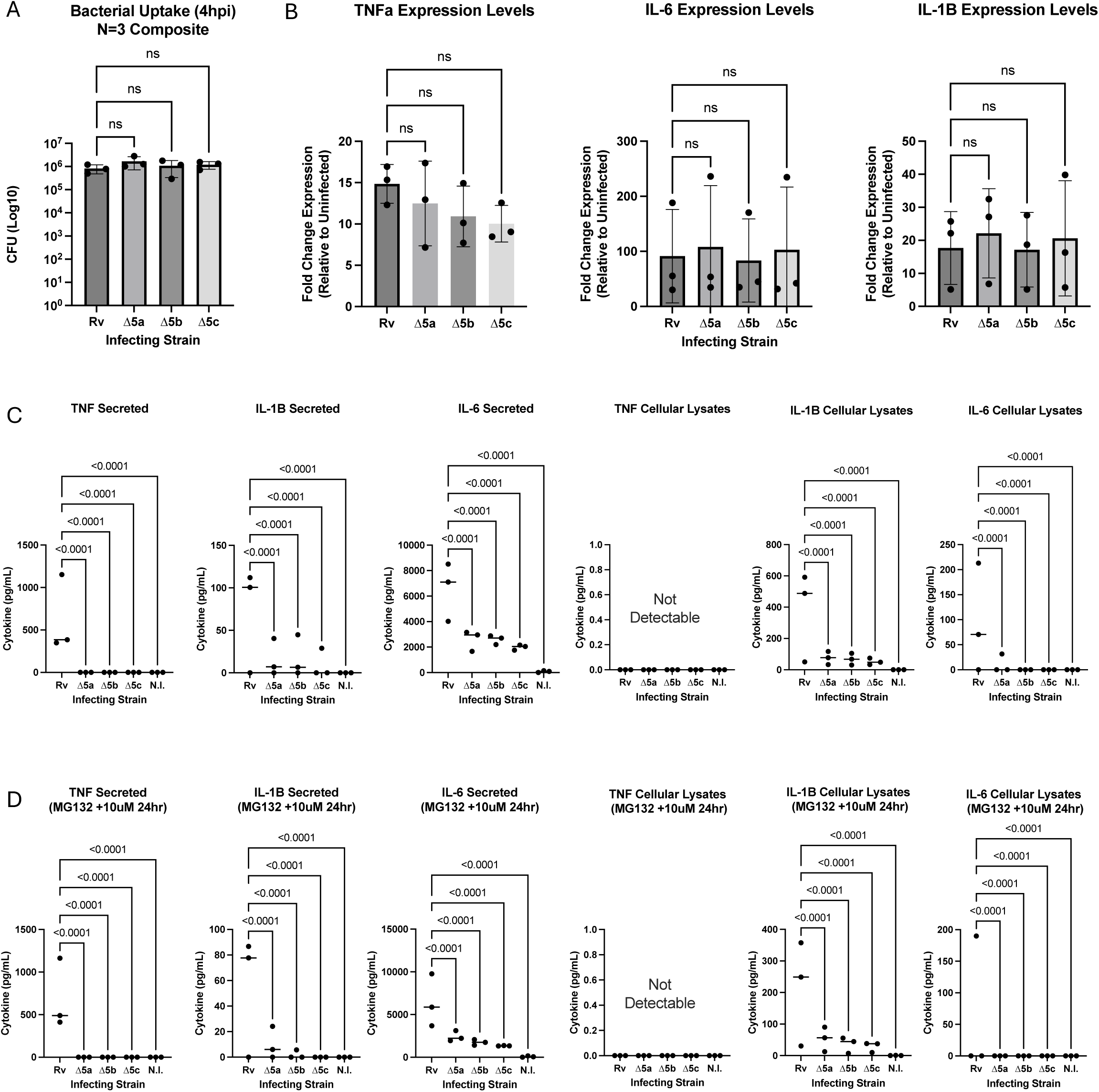
ESX-5 paralog deletions alter macrophage cytokine levels at post- transcriptional stage. *2A. Mtb macrophage uptake is unchanged across mutant strains*. Bar plot display 4-hour phagocytosis levels of respective strains (x-axis) measured via CFU (y-axis, log10 scale) for N=3 donors run in technical triplicate. *2B. Mtb induced cytokine mRNA induction is comparable across strains*. Bar charts display relative expression of TNF, IL-6, and IL-1β transcripts (y-axis, fold change expression) across N=3 donors (technical singlet) for respective strains (x-axis*). 2C. Mtb-induced cytokine protein is not retained with the cellular fraction of infected cells.* Dot plots represent TNF, IL-6, and IL-1β levels (y-axis pg/mL) for either secreted (Secreted) or cellular fractions (Cellular Lysates) of cells (N=3 donors, technical triplicate) infected with respective strains (x-axis, N.I = uninfected). *2D. Proteasome inhibition does not rescue cytokine protein levels.* Dot plots display paired graphs with 2C with additional treatment of 10µM MG132 (proteasome inhibition). *2A-2D.* Statistics generated using two-way ANOVA with post-hoc T-test and Dunnett correction for multiple comparisons (95% CI, p≤0.05). Error bars show standard deviation of the mean.

### Paralog deletions induce differential, target-specific regulation

To assess the breadth of the cytokine defect, we examined additional targets. Using a reporter assay we observed that IFNβ follows the same reduction trend as observed for the other cytokines (**Figure 3A**). In addition, CCL3, CCL5, and GM-CSF had reduced levels in ΔESX-5a, ΔESX-5b, and ΔESX-5c infected macrophages compared to wild type. In contrast, ΔESX-5a, ΔESX-5b, and ΔESX-5c induced comparable levels of IL-8 and CCL4 to wildtype Mtb (**Figure 3B, 3C**). Although IL-8 had high baseline levels that may obscured Mtb-dependent effects, CCL4 was highly induced and provided a contrast to the other cytokines. Together, these data suggest that ESX-5 deletions regulate Mtb-induced signaling pathways in a cytokine-specific manner.

**Figure 3.**
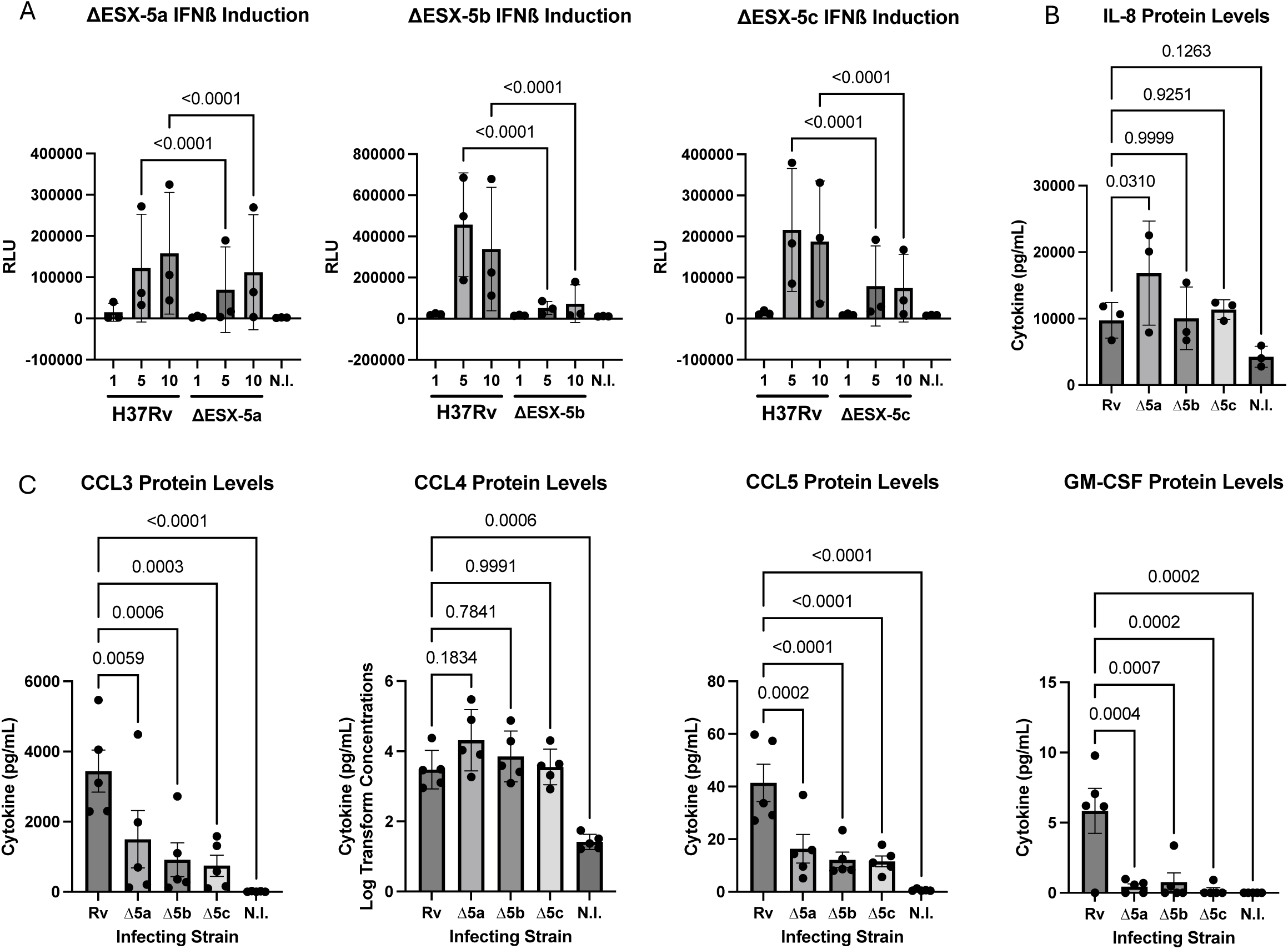
ESX-5 regulates Mtb-induced signaling in macrophages with cytokine specificity. *3A. Mtb-induced macrophage IFN-β is downregulated with mutant infection compared to wild type*. Bar charts display IFN-β response levels (y-axis, relative luminescence (RLU)) across N=3 donors (technical triplicate) and a bacterial dose curve (MOI 1, 5, or 10) comparing wildtype vs. respective knockout strains with an uninfected control (N.I.) (x-axis). *3B. IL-8 levels are unaffected by ESX-5 deletion mutants*. Bar chart shows the level of IL-8 expression (y-axis, pg/mL) across N=3 donors (technical triplicate) infected with respective strains (x-axis, N.I. = uninfected). *3C. Chemokines display heterogenous expression profile*. Bar charts display protein levels (y-axis, pg/mL) for four distinct chemokines (CCL3, CCL4, CCL5, and GM-CSF) across N=5 donors (technical singlet) for respective infection conditions (x-axis, N.I. = uninfected). CCL4 shows log transformed concentrations on y axis due to abnormal data across infection conditions (negative Shapiro-Wilk normality test). *3A-3C.* Statistics for all analyses generated using two-way ANOVA with post-hoc T-test and Dunnett correction for multiple comparisons (95% CI, p≤0.05). Error bars show standard deviation of the mean.

### ESX-5 knockouts induce similar miRNA profiles and cellular stress response relative to H37Rv infected MDMs

We next hypothesized that ESX-5 may regulate cytokine-specific secretion via small RNA-mediated translational inhibition. We examined micro-RNA (miRNA) profiles via nCounter miRNA profiling of macrophages infected with our strains (N=3 donors).

Among 827 biologically active and functional miRNA species, 130 were detectable (≥50 probes per sample threshold to remove low or inconsistently detected targets) **(Figure 4A, Supplemental Table 2)**. No miRNAs were differentially induced when comparing wild type to knockout infection conditions (p≤0.05). One probe displayed a marginal difference when comparing ΔESX-5c to wild type (hsa-miR-4454+hsa-miR-7975, p=0.1970) (**Figure 4B)**. We also queried whether ESX-5-dependent protein translation is regulated via the integrated stress response (ISR) or unfolded protein response (UPR)(50). Initially, we measured the levels of phospho-eIF2a as a measure of translation inhibition. Total detectable protein did not appear to differ within infected macrophages across strains but was highly variable (**Supplemental Figure 5**). Cells infected with mutant strains elicited similar levels of XBP1 splicing, HSPA5, and TRIB3 expression as markers of IRE1a, ATF6, and PERK pathway activation, respectively (**Figure 4C**). Using inhibitors of the UPR and ISR (4µ8c, Ceapin-A7, GSK2606414, and ISRIB targeting IRE1a, ATF6, PERK, and phosphorylation of eIF2a, respectively), we did not detect restoration of cytokine levels in ESX-5 mutant-infected macrophages for nearly all conditions tested (IL-6, TNF, or IL-1β) (**Supplemental Figure 6)**. Treatment with 4µ8c led to increased levels of IL-6 in H37Rv vs. ΔESX-5a infection; however, this was driven by 2/6 donors tested and was likely an outlier effect. Together, these analyses indicate that miRNA and UPR/ISR regulatory activation do not mediate the observed ESX-5 dependent post-transcriptional cytokine phenotype.

**Figure 4.**
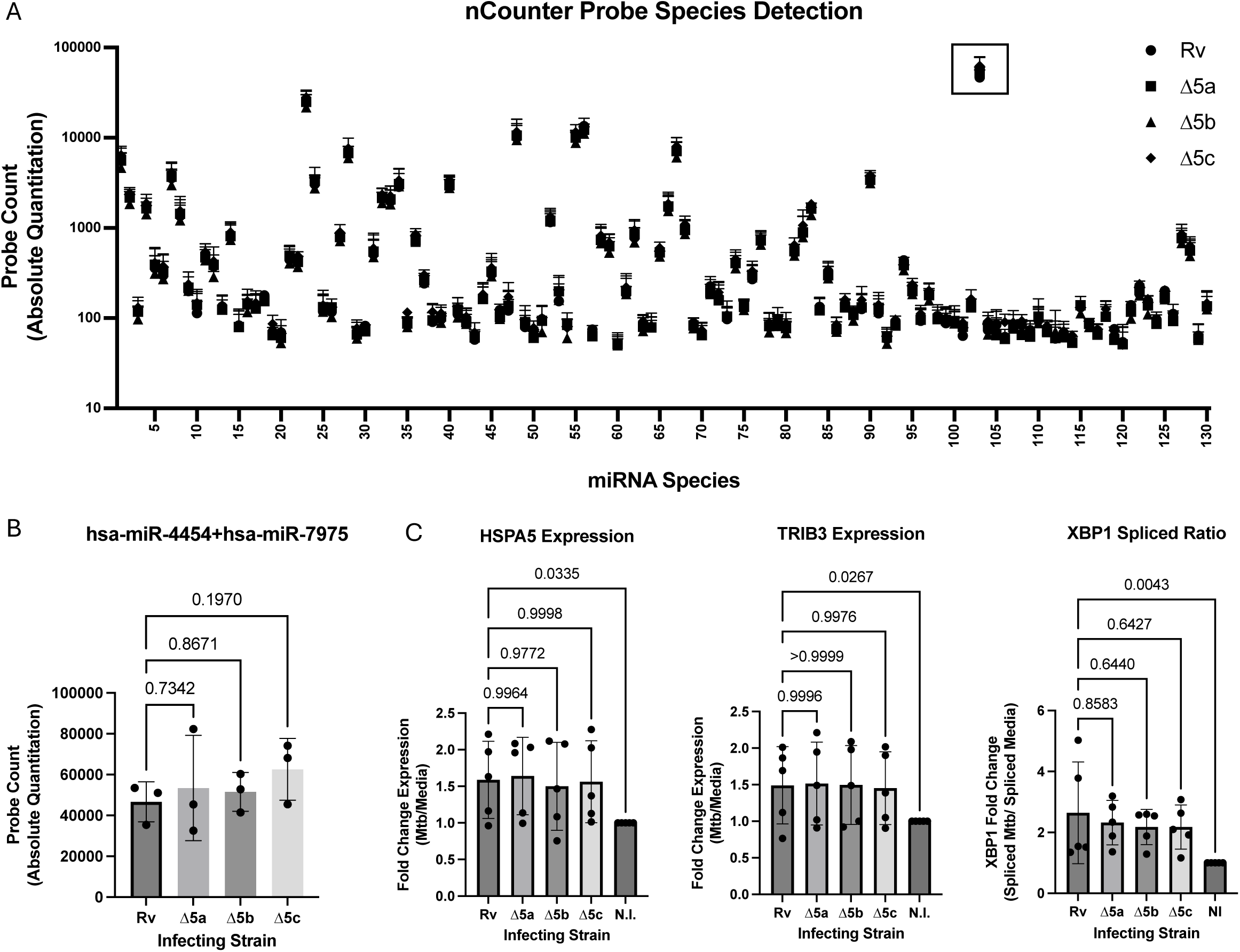
Macrophage miRNA and UPR responses are not ESX-5 dependent. *4A and B. Mtb-induced macrophage microRNA expression is not ESX-5-dependent.* Mtb inducible microRNA species (x-axis, number 1-130) in human macrophages (N=3, technical singlet) using nCounter (y-axis, absolute quantification) are shown. The included miRNA data has been baseline subtracted to remove low or inconsistently detected targets (removal of 697 probes, Dot shape correlates with respective strain (Circle = H37Rv, Square = Δ5a, Triangle = Δ5b, and Diamond = Δ5c). Black square shows highest abundance target for all samples (hsa-miR-4454+ hsa-miR-7975). 4B depicts microRNA hsa-miR-4454+ hsa-miR-7975 which had a marginal trend towards a difference between Δ5c and wild type. *4C. Mtb-induced macrophage UPR and ISR responses are not ESX-5-dependent*. UPR expression targets HSPA5, TRIB3, and spliced XBP1 are shown with fold change differences (y-axis, (Mtb/Media)) across respective strain infections (x-axis, N.I. = uninfected). Dots represent N=5 biological donors run in technical singlet for HSPA5 and TRIB3 while technical duplicate for XBP1. *4B-C*. Statistics generated using two-way ANOVA with post-hoc T-test and Dunnett correction for multiple comparisons (95% CI, p≤0.05). Error bars show standard deviation of the mean.

### Transcriptome sequencing of paralog knockouts reveals metal responsive gene signature

We next examined the transcriptional response of our ESX-5 knockouts in log phase growth to determine how knockout-dependent expression profiles in Mtb may impact macrophage cytokine responses. Compared to wild type, RNA sequencing revealed downregulation of 92, 88, and 85 genes in ΔESX-5a, ΔESX-5b, and ΔESX-5c, respectively (FDR ≤0.01, Log2 Fold-Change ≤-1, **Figure 5A, Supplemental Table 3-4)**. Seventy-one of these downregulated genes were shared between all three knockout strains and were used to generate a string network identifying connected clusters (**Figure 5B, Supplemental Figure 7, Supplemental Table 4).** Hypergeometric mean pathway analysis with Gene Ontology (GO) Biological Process revealed a strong and significant metal response gene signature, a function not previously associated with these gene clusters (**Figure 5C**). Specifically, “response to cadmium ion” (FDR 2.43E- 03) and “response to copper ion” (FDR 2.96E-03) were significantly enriched in our expression data (**Figure 5C**).

**Figure 5.**
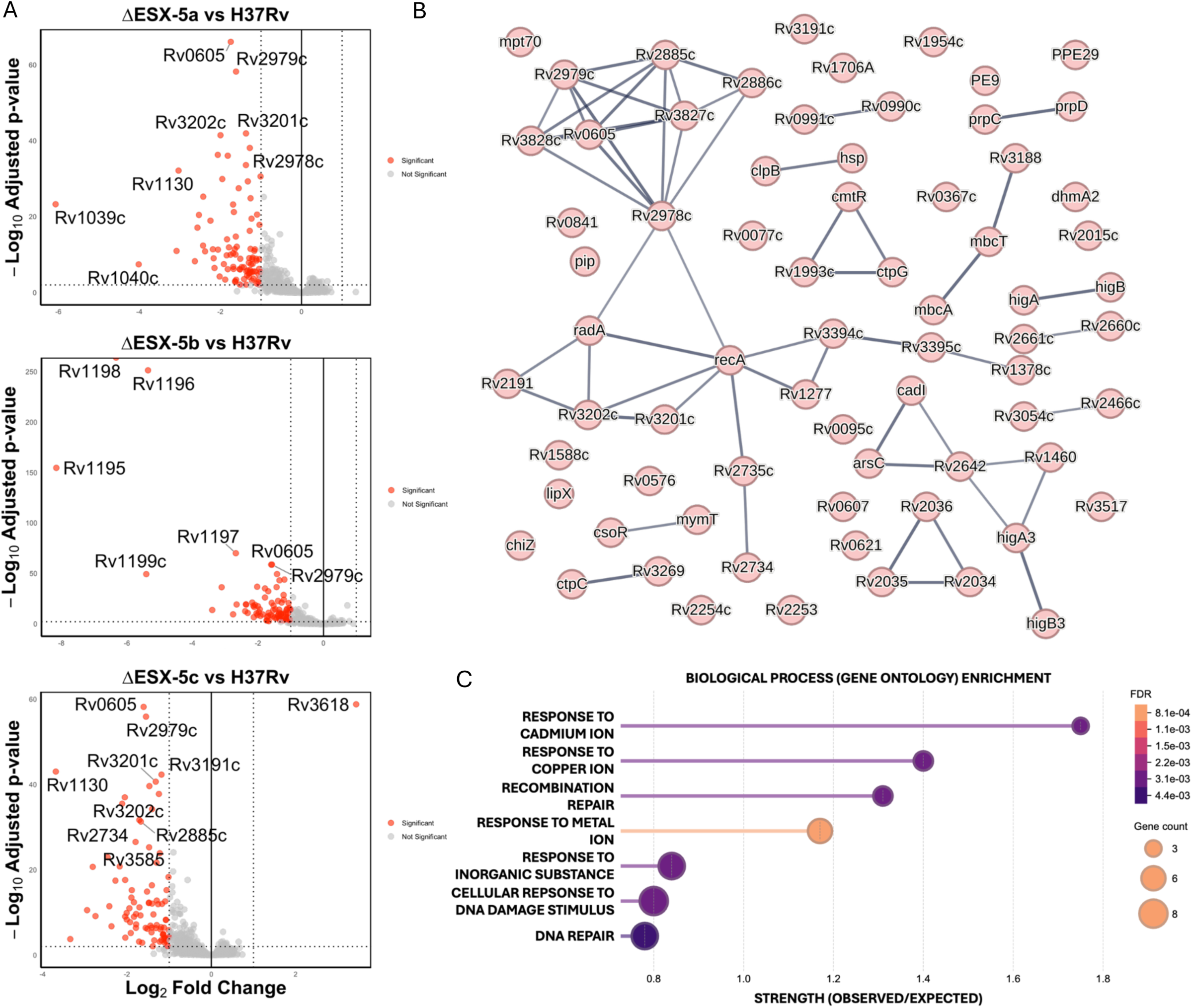
Bacterial transcriptional analysis of ESX-5 paralog knockouts reveals metal response axis. *5A. Volcano gene expression plots of mutants relative to wildtype.* Plots show respective expression profiles across mutant transcriptomes relative to wildtype. For all plots, significance level is on y-axis (Log10 FDR Corrected (“Adjusted”) p-value) and change in expression on x-axis (Log2 Fold Change). Red dots denote significant genes (FDR ≤0.01, Log2FC ≥1/≤-1), grey dots are non-significant. *5B. String network of overlapped down-regulated genes shared across all knockout strains*. Each node represents a gene while the connecting lines represent combined string score of respective nodes (minimum string score >0.700, medium high connectivity, string-db.org). *5C. Hypergeometric mean pathway enrichment analysis of overlapping genes.* Enrichment plot shows top 7 most significant gene sets (y-axis) based on Gene Ontology (GO) biological process. X-axis shows the strength of the observation (Log10(observed genes/expected genes)). Size of node proportional to the number of genes in our data belonging to a respective gene set. Color of node correlates to the FDR of the enrichment (significance calculated using Fishers Exact test with Benjamini- Hochberg FDR correction).

### Cadmium and copper exposure in Mtb causes increased ESX-5a and 5c transcription, increasing TNF and IL-6 production in macrophages

To assess whether ESX-5a, 5b, or 5c regulates Mtb metal responses, we treated strains with Cu, Cd, Fe, and Zn-supplemented 7H9 media over 5 days (100µM Cu/Cd or 500µM Fe/Zn). Cd and Cu restricted growth for all strains **(Figure 6A, Cd & Cu)**. In contrast, Fe and Zn treatment did not alter growth **(Figure 6A, Fe & Zn)**. There was no difference in growth between H37Rv or knockout strains, indicating no ESX-5- dependent sensitivity to these metals **(Figure 6A)**. To assess whether ESX-5a, 5b, and 5c are transcriptionally responsive to metal ions, we grew H37Rv in media containing high levels of Cd or Cu for 24 hours and analyzed the expression of ESX-5a, ESX-5b, and ESX-5c clusters via qPCR. Excess Cu upregulated ESX-5a and ESX-5c (**Figure 6B**). Conversely, Cu and Cd reduced ESX-5b expression, although only Cu reached significance (**Figure 6B).** To examine if there is a link between metal-induced ESX paralog expression and cytokine production, we pre-treated H37Rv Mtb for five days with either Cu or Cd in 7H9 broth, infected macrophages, and measured TNF, IL-1β, and IL-6. Pre-treatment of Mtb with these metals led to increased TNF and IL-6 (**Figure 6C**). Together, these data suggest that ESX-5 paralogs are associated with Mtb metal responses and that Cd/Cu regulate ESX-5a, 5b, and 5c gene expression while tuning cytokine responses in macrophage

**Figure 6.**
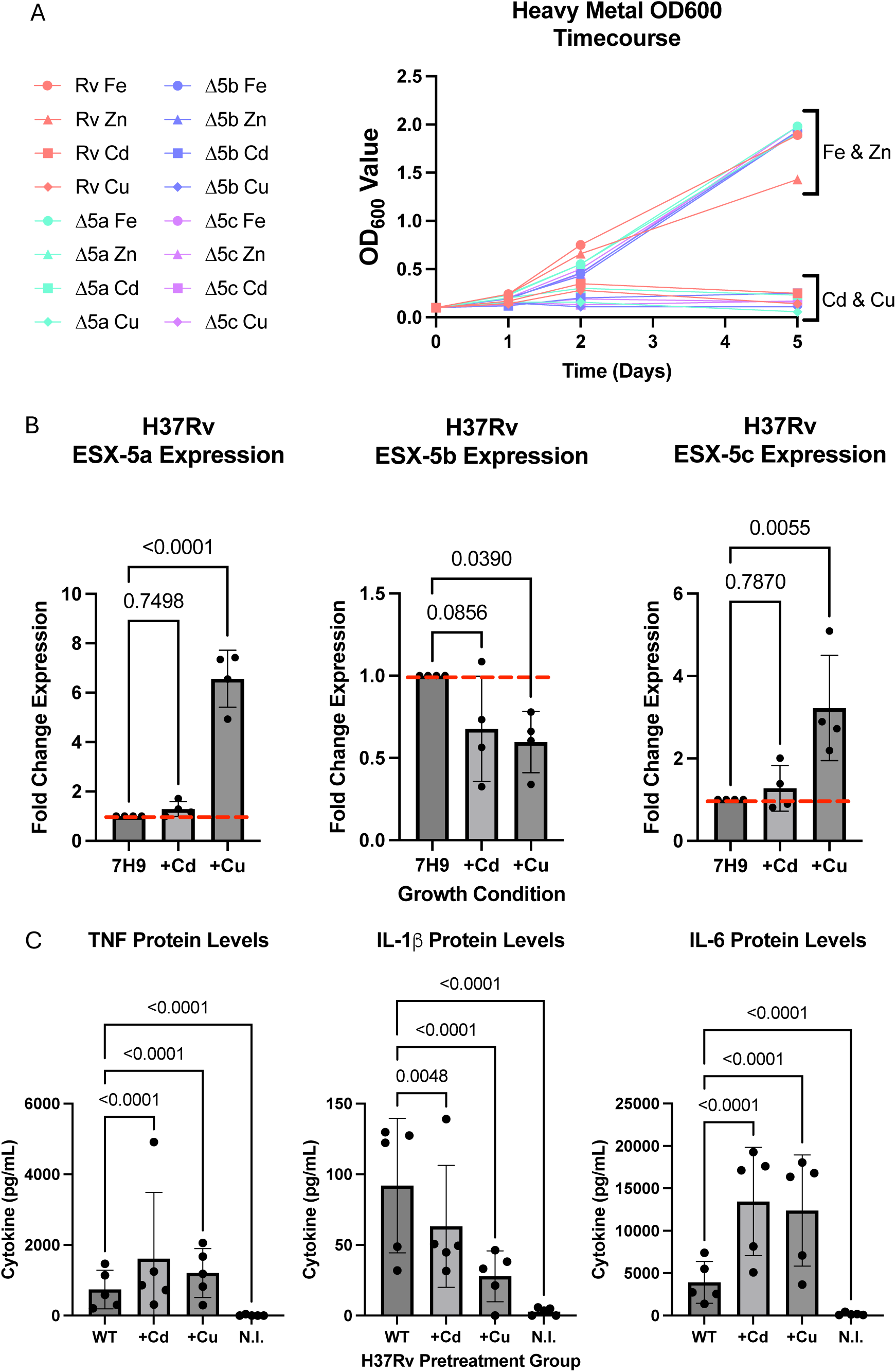
Bacterial heavy metal treatment leads to differential Mtb ESX-5 paralog expression and increased cytokine output in macrophages. ***6A. Mtb growth for*** *heavy metal treated strains*. Graph depicts 16 different growth curves (y-axis, OD600) (Paired strain (H37Rv = red, Δ5a = teal, Δ5b = blue, and Δ5c = pink) with respective metals (Iron 500µM = circle, Zinc 500µM = Triangle, Cadmium 100µM = Square, Copper 100µM = Diamond)) for 5 days (x-axis). Brackets on graph depict clustering of metal groups. *6B. Esx-5 paralog gene expression in H37Rv under metal stimulus*. Bar charts show expression (y-axis, fold change expression) of a respective Esx gene cluster (Title) within H37Rv under different stimulation conditions (x-axis, 7H9= untreated control) across N=4 replicate experiments. Red dotted line indicates baseline expression of respective gene cluster in untreated bacteria. *6C. Metal Mtb pre-treatment leads to increased cytokine expression in macrophages*. Bar charts show the relative abundance of TNF, IL-6, and IL-1β (y-axis, pg/mL) from N=5 donors (technical triplicate) infected with metal pre-treated H37Rv (x-axis, WT= no metal control, N.I. = uninfected). *6B-6C*. Statistics generated using two-way ANOVA with post-hoc T-test and Dunnett correction for multiple comparisons (95% CI, p≤0.05). Error bars show standard deviation of the mean.

## Discussion

Our primary findings demonstrate that ESX-5 regulates cytokine-specific macrophage responses via a post-transcriptional mechanism that is dependent on viable bacteria and not due to protein degradation or a block in protein secretion. Furthermore, ESX-5 regulates Mtb responses to metal, including Cd and Cu, which modulate macrophage cytokine secretion. Together, our data suggest that these three putative virulence clusters may play dual functions in regulating both macrophage inflammation and bacterial heavy metal responses.

Our *in vitro* data suggests a model whereby ESX-5-secreted molecules promote pro-inflammatory cellular responses as a mechanism for potentiating inflammation. Previous studies demonstrated that ESX-5 secreted proteins are inflammatory and immunogenic, generating potent T-cell responses and serving as candidate vaccine antigens (39,40). Our data suggest these effectors may also promote myeloid cell activation. Classically, human macrophages respond to Mtb infection by expressing proinflammatory cytokines including IL-6, TNF, and IL-1β (51–53). Although some pro-inflammatory responses have anti-microbial effects that benefit the host, there is also the potential that excessive pro-inflammatory responses may benefit Mtb. For example, inflammatory-induced sequelae can include cell death with release of bacilli and enhanced recruitment of myeloid cells to the site of infection (54–57). Our observations here suggest a model wherein ESX-5a, 5b, and 5c paralogs are actively secreted during early macrophage infection and promote inflammatory cytokine synthesis. This cellular state may alter the inflammatory disposition of host cells and in the context of human infection may promote enhanced leukocytic recruitment, immunopathology, and subsequent bacillary spread to naive immune cells.

The cellular mechanism of this ESX-5-dependent cytokine-specific phenotype suggests a post-transcriptional event. Our data demonstrate ESX-5 paralog knockout strains possess the correct stimulatory motifs to initiate inflammatory cascades with no differences in cytokine mRNA levels. Yet we observe a reduction in cytokine synthesis across divergent cytokine classes that are both cytosolically and ER translated.

Interestingly, the ESX-5 dependent protein levels are cytokine-specific with CCL4 and IL-8 (CXCL8) levels preserved during knockout infections. The regulatory motifs driving the expression of these cytokines relative to others is not fully understood, particularly in the context of Mtb infection. However, their unaltered expression levels suggests a post-transcriptional process that promotes their differential expression in contrast with the repression we observe for other cytokines such as IL-6 and TNF. Possible mechanisms governing this difference including transcript availability and initiation of protein translation. For example, TNF, IL-6 and other cytokine transcripts contain known RNA binding protein (RBP) motifs which recruit regulatory elements (58,59). Alternatively, some transcripts contain upstream ORF elements (uORF) which are ribosomal binding targets and could mediate cytokine-specific silencing (60). A potential model is that under wild type conditions, the ESX-5 effectors described here inhibit these regulatory elements and promote translation of inflammatory cytokines such as IL-6 and TNF. Conversely, during infection with the mutants these effectors are absent allowing for inhibitory regulation to occur within infected cells, effectively curbing synthesis of certain inflammatory markers. Whether these ESX-5 paralogs promote specific cytokine synthesis by directly inhibiting regulatory elements or indirectly via alternative mechanisms within the cell, remains to be elucidated.

Our data also demonstrates that ESX-5 paralogs regulate heavy metal responses, particularly to Cu and Cd. Previous studies indicate that some PE/PPE proteins mediate nutrient import and homeostasis of biologically critical ions including calcium and iron (33,61,62). Cu and Cd-induced ESX-5 responses could mediate several functions including metal transport for homeostatic balance, detection of specific microenvironments, or activation of bacterial stress/survival responses to metal exposure. To our knowledge, this is the first time ESX-5 secreted PE/PPE clusters have been implicated in metal homeostasis. A potential model could include sensing of heavy metal levels in the phagosome that differ from the extracellular environment and lead to differential regulation of the ESX-5 paralogs (63,64). This in turn could lead to the expression of ESX-5a, 5b, and 5c encoded PE/PPE heterodimers that stabilize metal-induced stress responses. In tandem, the encoded Esx heterodimers drive increased cytokine responses that may potentiate Mtb spread and survival, however this remains to be determined in the context of Mtb infection models. In support of this model, our ESX-5 complementation strains and metal pre-treated bacteria both upregulate these gene clusters and drive increased cytokine production in macrophages. Together, these data implicate a mechanism whereby heterodimeric pairs of proteins with distinct effector function are co-regulated to carry out independent tasks in a shared environment. This co-regulation of effectors is a common strategy used by many bacteria to link the expression of temporally or functionally related effectors (65).

Our study has several limitations. While primary human cells offer a physiologically relevant model for studying Mtb infection, donor variability can obscure some results. However, these consistent findings in the face of donor variability add rigor to our core observations and open possibilities for future mechanistic follow up in more experimentally tractable systems such as myeloid cell lines. An additional limitation is that we utilized a lab strain of Mtb (H37Rv) to perform these studies. Our use of the H37Rv strain, although advantageous due to its well-characterized genome and compatibility with existing reagents, may not fully capture the pathogenicity of more virulent or clinically relevant strains. Future studies with clinical isolates could enhance the translatability of our findings. Lastly, although we establish a strong correlation between metal availability and ESX-5 paralog expression, we have not directly demonstrated a mechanistic axis connecting metal levels with ESX-5 responses, broadly. Complementary approaches such as metal depletion assays and biophysical characterization of metal binding to our PE/PPE proteins of interest would help clarify these potential mechanisms. Together, these limitations highlight important directions for future research to dissect the cellular and molecular interactions between ESX-5 paralogs and the host.

In summary, we demonstrate that the ESX-5 paralogs, ESX-5a, ESX-5b, and ESX-5c, play multiple roles in tuning inflammatory responses in macrophages via post-transcriptional pathways while also regulating Mtb responses to metal stimulation. Our study highlights the importance of investigating Mtb genes in the context of bacterial and cellular responses to gain more complete insight into gene function across diverse environments.

## Declarations

### Data and Materials Availability

Informatics tools and workflows used within this study are available on GitHub (https://github.com/BIGslu/SEAsnake,https://github.com/BIGslu/tutorials/tree/main/RNA <u>seq</u>). R software for performing subsequent analyses is available via the CRAN repository (https://cran.r-project.org/). All additional data has been published with this study or is available from the authors upon reasonable request. Biological materials generated within this study will be shared upon request in accordance with all applicable laws and regulations.

## Author Contributions

Study Conception: A.M.H, T.R.H, J.S.C

Experimental Procedures: A.M.H, S.B.C, A.P

Sample Processing: A.M.H, S.B.C, A.P

Data Analysis: A.M.H, S.B.C, T.R.H, J.S.C, D.R.S, K.B.U

Manuscript Drafting: A.M.H, T.R.H

Manuscript Editing: All Authors

## Acknowledgements

We thank the many individuals who contributed to make this work possible. Drs. Christoph Grundner and Andrew Frando for their gift of the H37Rv pNIT strain and technical advice deploying recombineering approaches. Dr. Nathan Keiswetter and Jessica Assadi for their technical assistance running flow cytometry and instruction on Mtb library preparation for sequencing, respectively. Drs. Kim Dill-McFarland and Max Segnitz for informatics analysis advice and support. Glenna Peterson for all primary human PBMC isolations and cell stock generation. And lastly Moeko Agata for her help in performing ELISA.

## References

1. Gagneux S. Ecology and evolution of Mycobacterium tuberculosis. Nat Rev Microbiol. 2018 Apr;16(4):202–13.

2. Pai M, Behr MA, Dowdy D, Dheda K, Divangahi M, Boehme CC, et al. Tuberculosis. Nat Rev Dis Primer. 2016 Dec 22;2(1):16076.

3. Goletti D, Meintjes G, Andrade BB, Zumla A, Lee SS. Insights from the 2024 WHO Global Tuberculosis Report – More Comprehensive Action, Innovation, and Investments required for achieving WHO End TB goals. Int J Infect Dis [Internet]. 2025 Jan 1 [cited 2025 May 1];150. Available from: https://www.ijidonline.com/article/S1201-9712%2824%2900400-4/fulltext

4. Glaziou P, Sismanidis C, Floyd K, Raviglione M. Global Epidemiology of Tuberculosis. Cold Spring Harb Perspect Med. 2015 Feb;5(2):a017798.

5. Global Tuberculosis Report 2024. 1st ed. Geneva: World Health Organization; 2024. 1 p.

6. McHenry ML, Bartlett J, Igo RP, Wampande EM, Benchek P, Mayanja-Kizza H, et al. Interaction between host genes and Mycobacterium tuberculosis lineage can affect tuberculosis severity: Evidence for coevolution? Schurr E, editor. PLOS Genet. 2020 Apr 30;16(4):e1008728.

7. Chandra P, Grigsby SJ, Philips JA. Immune evasion and provocation by Mycobacterium tuberculosis. Nat Rev Microbiol. 2022 Dec;20(12):750–66.

8. Silva Miranda M, Breiman A, Allain S, Deknuydt F, Altare F. The Tuberculous Granuloma: An Unsuccessful Host Defence Mechanism Providing a Safety Shelter for the Bacteria? Clin Dev Immunol. 2012;2012:139127.

9. Flynn JL, Chan J, Lin PL. Macrophages and control of granulomatous inflammation in tuberculosis. Mucosal Immunol. 2011 May;4(3):271–8.

10. Wong D, Bach H, Sun J, Hmama Z, Av-Gay Y. *Mycobacterium tuberculosis* protein tyrosine phosphatase (PtpA) excludes host vacuolar-H ^+^ –ATPase to inhibit phagosome acidification. Proc Natl Acad Sci. 2011 Nov 29;108(48):19371–6.

11. Fernandez-Soto P, Bruce AJE, Fielding AJ, Cavet JS, Tabernero L. Mechanism of catalysis and inhibition of Mycobacterium tuberculosis SapM, implications for the development of novel antivirulence drugs. Sci Rep. 2019 Dec;9(1):10315.

12. Chai Q, Yu S, Zhong Y, Lu Z, Qiu C, Yu Y, et al. A bacterial phospholipid phosphatase inhibits host pyroptosis by hijacking ubiquitin. Science. 2022 Oct 14;378(6616):eabq0132.

13. Forrellad MA, Klepp LI, Gioffré A, Sabio y García J, Morbidoni HR, Santangelo M de la P, et al. Virulence factors of the Mycobacterium tuberculosis complex. Virulence. 2013 Jan 1;4(1):3–66.

14. Feltcher ME, Sullivan JT, Braunstein M. Protein export systems of Mycobacterium tuberculosis: novel targets for drug development? Future Microbiol. 2010 Oct;5:1581– 97.

15. Miller BK, Zulauf KE, Braunstein M. The Sec Pathways and Exportomes of Mycobacterium tuberculosis. Microbiol Spectr. 2017 Apr 7;5(2):10.1128/microbiolspec.tbtb2-0013–2016.

16. Rivera-Calzada A, Famelis N, Llorca O, Geibel S. Type VII secretion systems: structure, functions and transport models. Nat Rev Microbiol. 2021 Sept;19(9):567–84.

17. Bunduc CM, Fahrenkamp D, Wald J, Ummels R, Bitter W, Houben ENG, et al. Structure and dynamics of a mycobacterial type VII secretion system. Nature. 2021 May 20;593(7859):445–8.

18. Cole ST, Brosch R, Parkhill J, Garnier T, Churcher C, Harris D, et al. Deciphering the biology of Mycobacterium tuberculosis from the complete genome sequence. Nature. 1998 June;393(6685):537–44.

19. Famelis N, Geibel S, Tol D van. Mycobacterial type VII secretion systems. Biol Chem. 2023 June 1;404(7):691–702.

20. Stoop EJM, Bitter W, Sar AM van der. Tubercle bacilli rely on a type VII army for pathogenicity. Trends Microbiol. 2012 Oct 1;20(10):477–84.

21. Chen X, Cheng H fu, Zhou J, Chan C Yeung, Lau K Fai, Tsui SK Wing, et al. Structural basis of the PE–PPE protein interaction in Mycobacterium tuberculosis. J Biol Chem. 2017 Oct 13;292(41):16880–90.

22. Pathak SK, Basu S, Basu KK, Banerjee A, Pathak S, Bhattacharyya A, et al. Direct extracellular interaction between the early secreted antigen ESAT-6 of Mycobacterium tuberculosis and TLR2 inhibits TLR signaling in macrophages. Nat Immunol. 2007 June;8(6):610–8.

23. Osman MM, Shanahan JK, Chu F, Takaki KK, Pinckert ML, Pagán AJ, et al. The C terminus of the mycobacterium ESX-1 secretion system substrate ESAT-6 is required for phagosomal membrane damage and virulence. Proc Natl Acad Sci. 2022 Mar 15;119(11):e2122161119.

24. Stanley SA, Johndrow JE, Manzanillo P, Cox JS. The Type I IFN Response to Infection with *Mycobacterium tuberculosis* Requires ESX-1-Mediated Secretion and Contributes to Pathogenesis. J Immunol. 2007 Mar 1;178(5):3143–52.

25. Abdallah AM, Verboom T, Weerdenburg EM, Gey van Pittius NC, Mahasha PW, Jiménez C, et al. PPE and PE_PGRS proteins of *Mycobacterium marinum* are transported via the type VII secretion system ESX-5. Mol Microbiol. 2009 Aug;73(3):329–40.

26. Bunduc CM, Ding Y, Kuijl C, Marlovits TC, Bitter W, Houben ENG. Reconstitution of a minimal ESX-5 type VII secretion system suggests a role for PPE proteins in the outer membrane transport of proteins. Ellermeier CD, editor. mSphere. 2023 Oct 24;8(5):e00402–23.

27. De Maio F, Berisio R, Manganelli R, Delogu G. PE_PGRS proteins of Mycobacterium tuberculosis: A specialized molecular task force at the forefront of host–pathogen interaction. Virulence. 11(1):898–915.

28. Ates LS. New insights into the mycobacterial PE and PPE proteins provide a framework for future research. Mol Microbiol. 2020 Jan;113(1):4–21.

29. Ates LS, Dippenaar A, Ummels R, Piersma SR, Van Der Woude AD, Van Der Kuij K, et al. Mutations in ppe38 block PE_PGRS secretion and increase virulence of Mycobacterium tuberculosis. Nat Microbiol. 2018 Jan 15;3(2):181–8.

30. Ates LS, Sayes F, Frigui W, Ummels R, Damen MPM, Bottai D, et al. RD5-mediated lack of PE_PGRS and PPE-MPTR export in BCG vaccine strains results in strong reduction of antigenic repertoire but little impact on protection. PLOS Pathog. 2018 June 18;14(6):e1007139.

31. Izquierdo Lafuente B, Ummels R, Kuijl C, Bitter W, Speer A. Mycobacterium tuberculosis Toxin CpnT Is an ESX-5 Substrate and Requires Three Type VII Secretion Systems for Intracellular Secretion. mBio. 2021 Apr 27;12(2):e02983–20.

32. Ates LS, van der Woude AD, Bestebroer J, van Stempvoort G, Musters RJP, Garcia- Vallejo JJ, et al. The ESX-5 System of Pathogenic Mycobacteria Is Involved In Capsule Integrity and Virulence through Its Substrate PPE10. Behr MA, editor. PLOS Pathog. 2016 June 9;12(6):e1005696.

33. Wang Q, Boshoff HIM, Harrison JR, Ray PC, Green SR, Wyatt PG, et al. PE/PPE proteins mediate nutrient transport across the outer membrane of Mycobacterium tuberculosis. Science. 2020 Mar 6;367(6482):1147–51.

34. Gröschel MI, Sayes F, Simeone R, Majlessi L, Brosch R. ESX secretion systems: mycobacterial evolution to counter host immunity. Nat Rev Microbiol. 2016 Nov;14(11):677–91.

35. Dong D, Wang D, Li M, Wang H, Yu J, Wang C, et al. PPE38 Modulates the Innate Immune Response and Is Required for Mycobacterium marinum Virulence. Flynn JL, editor. Infect Immun. 2012 Jan;80(1):43–54.

36. Shah S, Cannon JR, Fenselau C, Briken V. A Duplicated ESAT-6 Region of ESX-5 Is Involved in Protein Export and Virulence of Mycobacteria. Infect Immun. 2015 Nov;83(11):4349–61.

37. Holt KE, McAdam P, Thai PVK, Thuong NTT, Ha DTM, Lan NN, et al. Frequent transmission of the Mycobacterium tuberculosis Beijing lineage and positive selection for the EsxW Beijing variant in Vietnam. Nat Genet. 2018 June;50(6):849–56.

38. Saelens JW, Sweeney MI, Viswanathan G, Xet-Mull AM, Jurcic Smith KL, Sisk DM, et al. An ancestral mycobacterial effector promotes dissemination of infection. Cell. 2022 Nov;S0092867422013617.

39. Stylianou E, Pinpathomrat N, Sampson O, Richard A, Korompis M, McShane H. A five- antigen Esx-5a fusion delivered as a prime-boost regimen protects against M.tb challenge. Front Immunol. 2023 Oct 5;14:1263457.

40. Stylianou E, Harrington-Kandt R, Beglov J, Bull N, Pinpathomrat N, Swarbrick GM, et al. Identification and Evaluation of Novel Protective Antigens for the Development of a Candidate Tuberculosis Subunit Vaccine. Ehrt S, editor. Infect Immun. 2018 July;86(7):e00014–18.

41. Murphy KC, Papavinasasundaram K, Sassetti CM. Mycobacterial Recombineering. In: Parish T, Roberts DM, editors. Mycobacteria Protocols [Internet]. New York, NY: Springer New York; 2015 [cited 2023 May 4]. p. 177–99. (Methods in Molecular Biology; vol. 1285). Available from: https://link.springer.com/10.1007/978-1-4939-2450-9_10

42. Rock JM, Hopkins FF, Chavez A, Diallo M, Chase MR, Gerrick ER, et al. Programmable transcriptional repression in mycobacteria using an orthogonal CRISPR interference platform. Nat Microbiol. 2017 Apr;2(4):16274.

43. Fuss IJ, Kanof ME, Smith PD, Zola H. Isolation of whole mononuclear cells from peripheral blood and cord blood. Curr Protoc Immunol. 2009 Apr;Chapter 7:7.1.1-7.1.8.

44. Smith JR, Dowling JW, McFadden MI, Karp A, Schwerk J, Woodward JJ, et al. MEF2A suppresses stress responses that trigger DDX41-dependent IFN production. Cell Rep. 2023 Aug;42(8):112805.

45. Ortiz-Lazareno PC, Hernandez-Flores G, Dominguez-Rodriguez JR, Lerma-Diaz JM, Jave-Suarez LF, Aguilar-Lemarroy A, et al. MG132 proteasome inhibitor modulates proinflammatory cytokines production and expression of their receptors in U937 cells: involvement of nuclear factor-κB and activator protein-1. Immunology. 2008 Aug;124(4):534–41.

46. Culviner PH, Guegler CK, Laub MT. A Simple, Cost-Effective, and Robust Method for rRNA Depletion in RNA-Sequencing Studies. Cooper VS, editor. mBio. 2020 Apr 28;11(2):e00010–20.

47. Dill-McFarland K, Benson B, rmsegnitz. BIGslu/SEAsnake: v1.1 [Internet]. Zenodo; 2024 [cited 2025 June 22]. Available from: https://zenodo.org/records/11646755

48. Michael Love SA. DESeq2 [Internet]. Bioconductor; 2017 [cited 2025 June 26]. Available from: https://bioconductor.org/packages/DESeq2

49. Di Luca M, Bottai D, Batoni G, Orgeur M, Aulicino A, Counoupas C, et al. The ESX-5 Associated eccB5-eccC5 Locus Is Essential for Mycobacterium tuberculosis Viability. Manganelli R, editor. PLoS ONE. 2012 Dec 17;7(12):e52059.

50. Xu P, Tang J, He ZG. Induction of Endoplasmic Reticulum Stress by CdhM Mediates Apoptosis of Macrophage During Mycobacterium tuberculosis Infection. Front Cell Infect Microbiol. 2022 Apr 4;12:877265.

51. Marino S, Sud D, Plessner H, Lin PL, Chan J, Flynn JL, et al. Differences in Reactivation of Tuberculosis Induced from Anti-TNF Treatments Are Based on Bioavailability in Granulomatous Tissue. Wodarz D, editor. PLoS Comput Biol. 2007 Oct 19;3(10):e194.

52. Martinez AN, Mehra S, Kaushal D. Role of Interleukin 6 in Innate Immunity to Mycobacterium tuberculosis Infection. J Infect Dis. 2013 Apr 15;207(8):1253–61.

53. Silvério D, Gonçalves R, Appelberg R, Saraiva M. Advances on the Role and Applications of Interleukin-1 in Tuberculosis. mBio. 12(6):e03134–21.

54. Beckwith KS, Beckwith MS, Ullmann S, Sætra RS, Kim H, Marstad A, et al. Plasma membrane damage causes NLRP3 activation and pyroptosis during Mycobacterium tuberculosis infection. Nat Commun. 2020 May 8;11(1):2270.

55. Abdallah AM, Bestebroer J, Savage NDL, de Punder K, van Zon M, Wilson L, et al. Mycobacterial Secretion Systems ESX-1 and ESX-5 Play Distinct Roles in Host Cell Death and Inflammasome Activation. J Immunol. 2011 Nov 1;187(9):4744–53.

56. Afriyie-Asante A, Dabla A, Dagenais A, Berton S, Smyth R, Sun J. Mycobacterium tuberculosis Exploits Focal Adhesion Kinase to Induce Necrotic Cell Death and Inhibit Reactive Oxygen Species Production. Front Immunol. 2021 Oct 20;12:742370.

57. Butler RE, Brodin P, Jang J, Jang MS, Robertson BD, Gicquel B, et al. The Balance of Apoptotic and Necrotic Cell Death in Mycobacterium tuberculosis Infected Macrophages Is Not Dependent on Bacterial Virulence. Briken V, editor. PLoS ONE. 2012 Oct 30;7(10):e47573.

58. Cook ME, Bradstreet TR, Webber AM, Kim J, Santeford A, Harris KM, et al. The ZFP36 family of RNA binding proteins regulates homeostatic and autoreactive T cell responses. Sci Immunol. 2022 Oct 28;7(76):eabo0981.

59. Otsuka H, Fukao A, Funakami Y, Duncan KE, Fujiwara T. Emerging Evidence of Translational Control by AU-Rich Element-Binding Proteins. Front Genet [Internet]. 2019 May 2 [cited 2025 July 21];10. Available from: https://www.frontiersin.org/journals/genetics/articles/10.3389/fgene.2019.00332/full

60. Zhong Z, Li Y, Sun Q, Chen D. Tiny but mighty: Diverse functions of uORFs that regulate gene expression. Comput Struct Biotechnol J. 2024 Dec;23:3771–9.

61. Boradia V, Frando A, Grundner C. The Mycobacterium tuberculosis PE15/PPE20 complex transports calcium across the outer membrane. PLOS Biol. 2022 Nov 28;20(11):e3001906.

62. Tufariello JM, Chapman JR, Kerantzas CA, Wong KW, Vilchèze C, Jones CM, et al. Separable roles for Mycobacterium tuberculosis ESX-3 effectors in iron acquisition and virulence. Proc Natl Acad Sci U S A. 2016 Jan 19;113(3):E348–57.

63. Sheldon JR, Skaar EP. Metals as phagocyte antimicrobial effectors. Curr Opin Immunol. 2019 Oct;60:1–9.

64. Samanovic MI, Ding C, Thiele DJ, Darwin KH. Copper in microbial pathogenesis: meddling with the metal. Cell Host Microbe. 2012 Feb 16;11(2):106–15.

65. Jagadeesan R, Dash S, Palma CSD, Baptista ISC, Chauhan V, Mäkelä J, et al. Dynamics of bacterial operons during genome-wide stresses is influenced by premature terminations and internal promoters. Sci Adv. 2025 May 16;11(20):eadl3570.

