## Supplemental Figures and Legends for "ESX-5 Deletions in *Mycobacterium tuberculosis* Alter Macrophage Cytokine Signaling and Bacterial Heavy Metal Response"

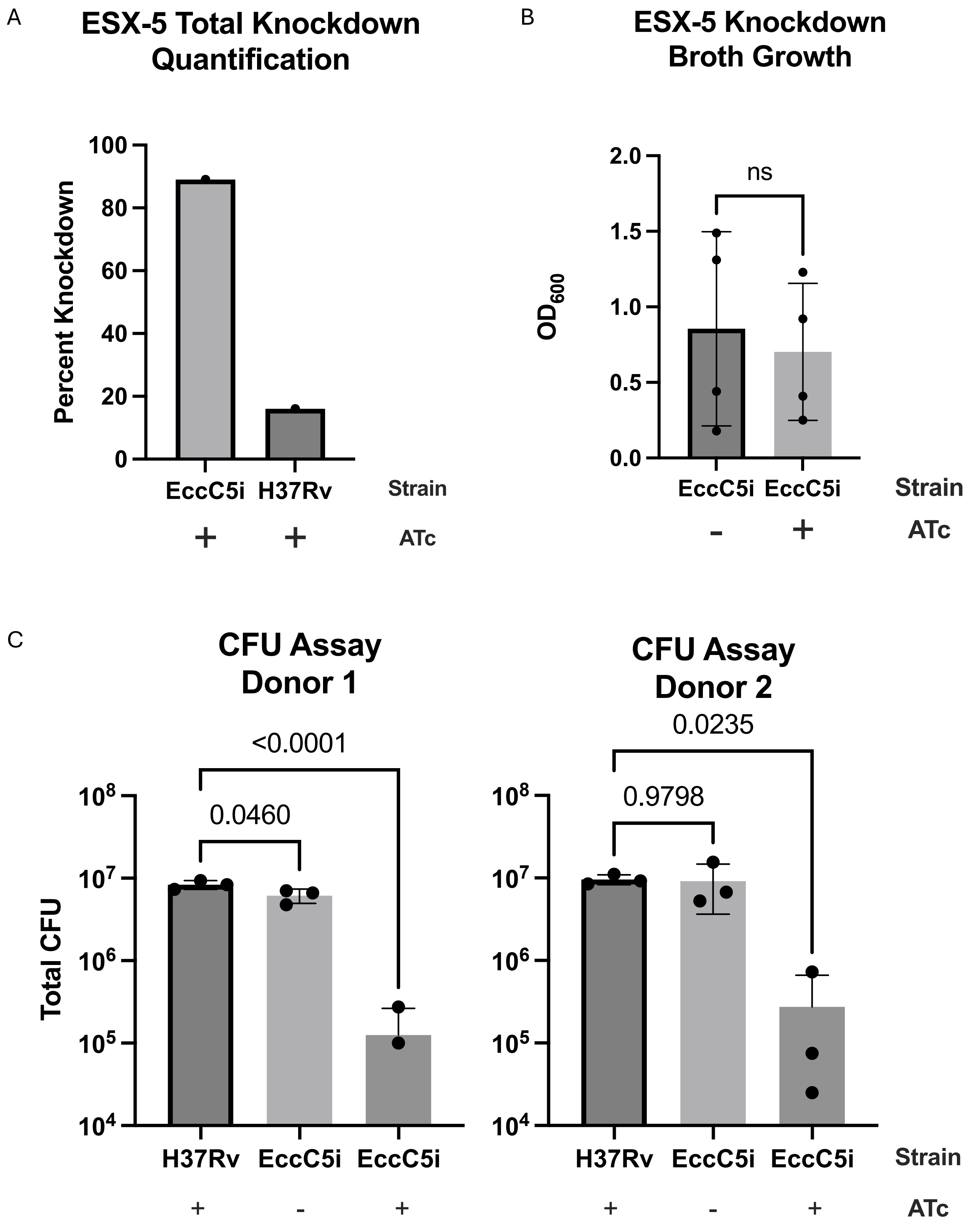
A
ESX-5 Total Knockdown Quantification
Percent Knockdown
0
20
40
60
80
100
EccC5i
H37Rv
Strain
+
+
ATc
B
ESX-5 Knockdown Broth Growth
OD600
0.0
0.5
1.0
1.5
2.0
ns
EccC5i
EccC5i
Strain
-
+
ATc
C
CFU Assay Donor 1
Total CFU
10^4
10^5
10^6
10^7
10^8
0.0460
<0.0001
H37Rv
EccC5i
EccC5i
+
-
+
CFU Assay Donor 2
10^4
10^5
10^6
10^7
10^8
0.9798
0.0235
H37Rv
EccC5i
EccC5i
Strain
+
-
+
ATc

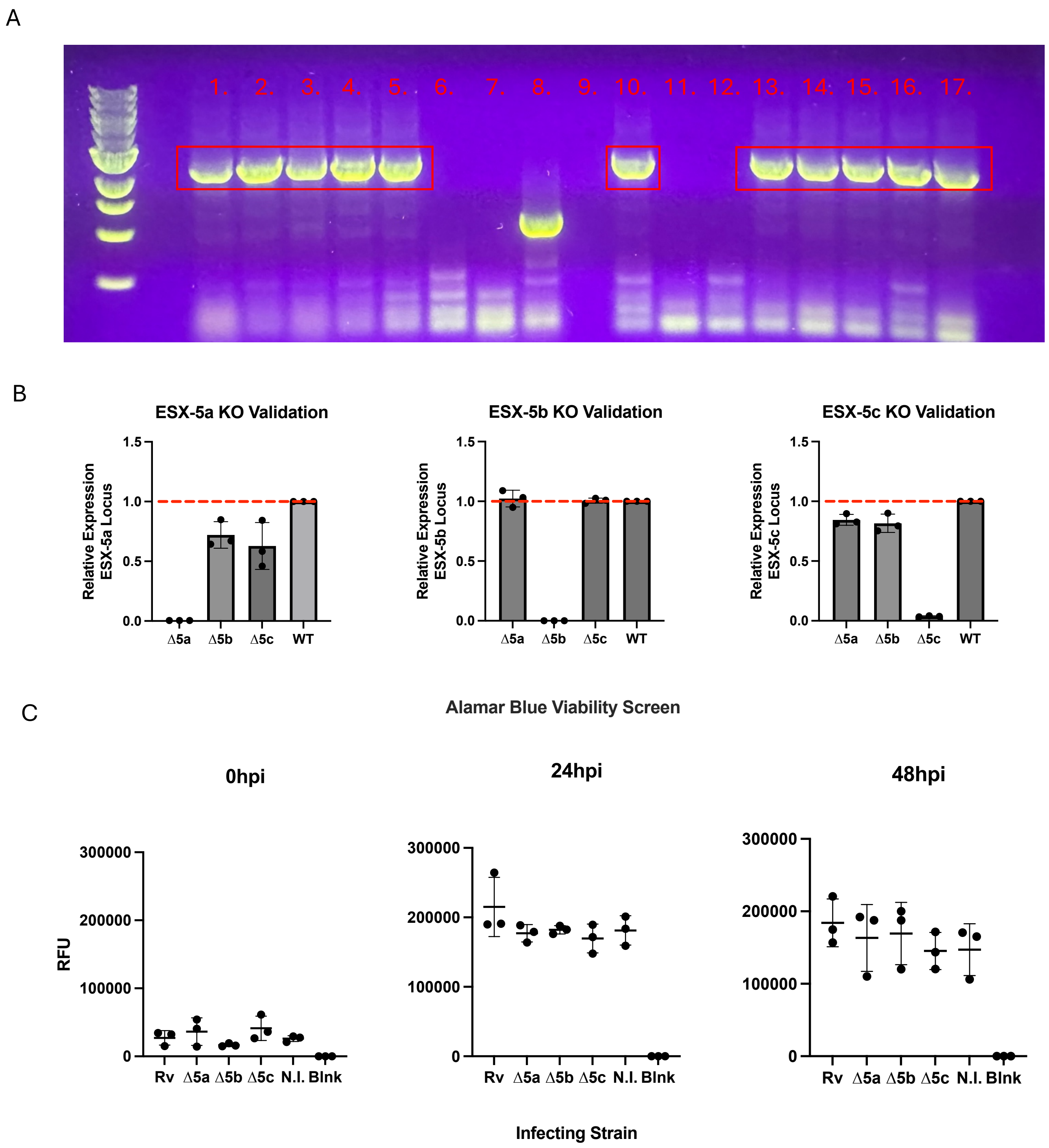
A
1. 2. 3. 4. 5. 6. 7. 8. 9. 10. 11. 12. 13. 14. 15. 16. 17.
B
ESX-5a KO Validation
Relative Expression ESX-5a Locus
1.5
1.0
0.5
0.0
Δ5a Δ5b Δ5c WT
ESX-5b KO Validation
Relative Expression ESX-5b Locus
Δ5a Δ5b Δ5c WT
ESX-5c KO Validation
Relative Expression ESX-5c Locus
Δ5a Δ5b Δ5c WT
C
Alamar Blue Viability Screen
0hpi
24hpi
48hpi
RFU
300000
200000
100000
0
Rv Δ5a Δ5b Δ5c N.I. Blnk
Infecting Strain

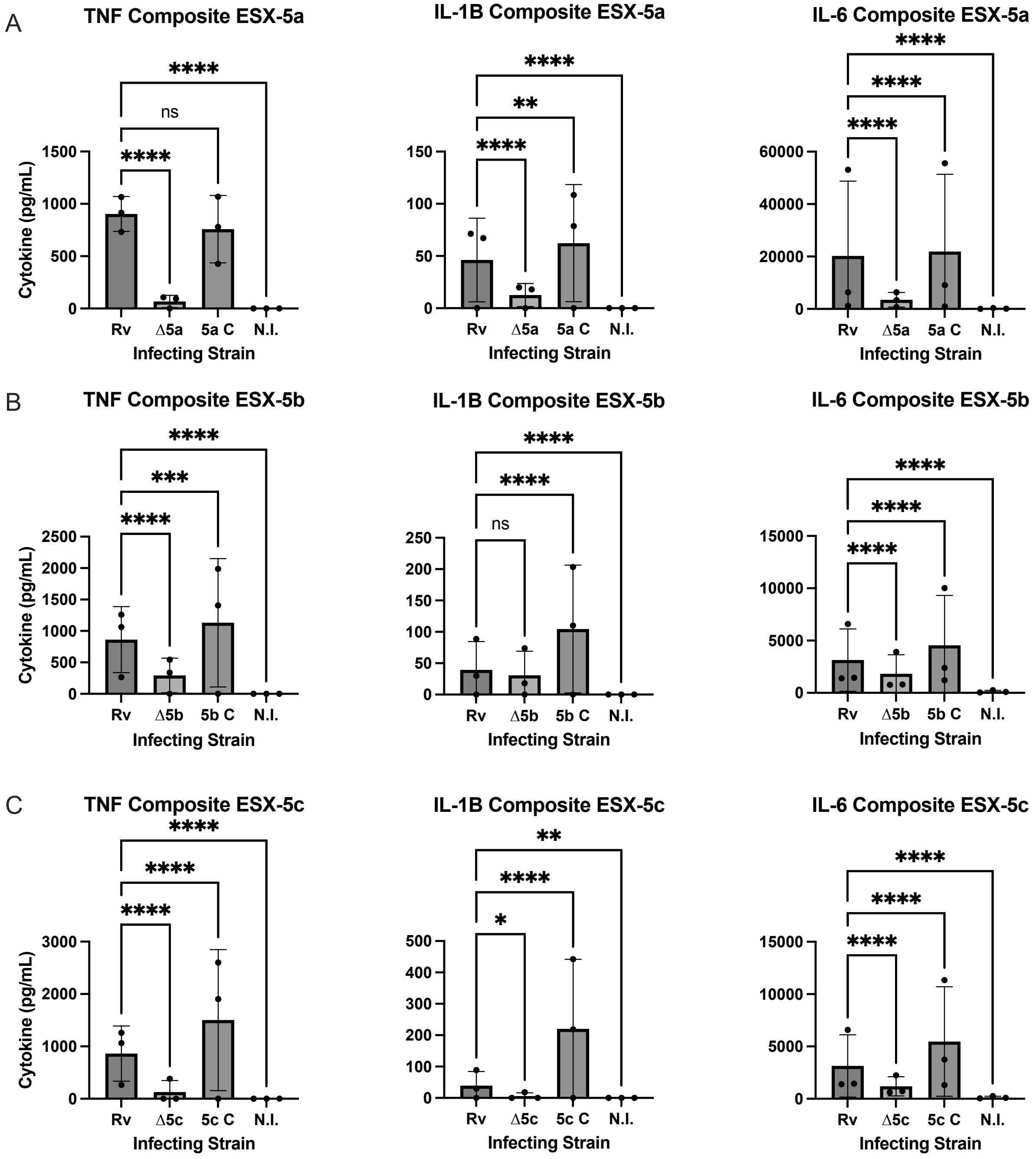

A
TNF Composite ESX-5a
IL-1B Composite ESX-5a
IL-6 Composite ESX-5a
B
TNF Composite ESX-5b
IL-1B Composite ESX-5b
IL-6 Composite ESX-5b
C
TNF Composite ESX-5c
IL-1B Composite ESX-5c
IL-6 Composite ESX-5c
Cytokine (pg/mL)
Infecting Strain
Rv
Δ5a
5a C
Δ5b
5b C
Δ5c
5c C
N.I.
ns

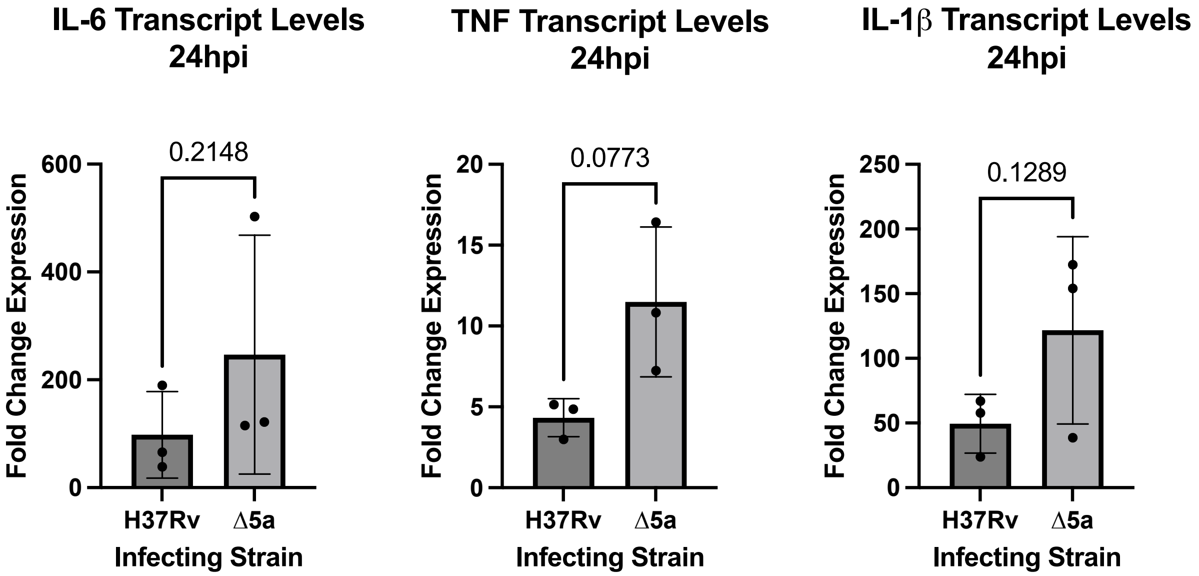
IL-6 Transcript Levels
24hpi
0.2148
Fold Change Expression
600
400
200
0
H37Rv
Δ5a
Infecting Strain
TNF Transcript Levels
24hpi
0.0773
Fold Change Expression
20
15
10
5
0
H37Rv
Δ5a
Infecting Strain
IL-1β Transcript Levels
24hpi
0.1289
Fold Change Expression
250
200
150
100
50
0
H37Rv
Δ5a
Infecting Strain

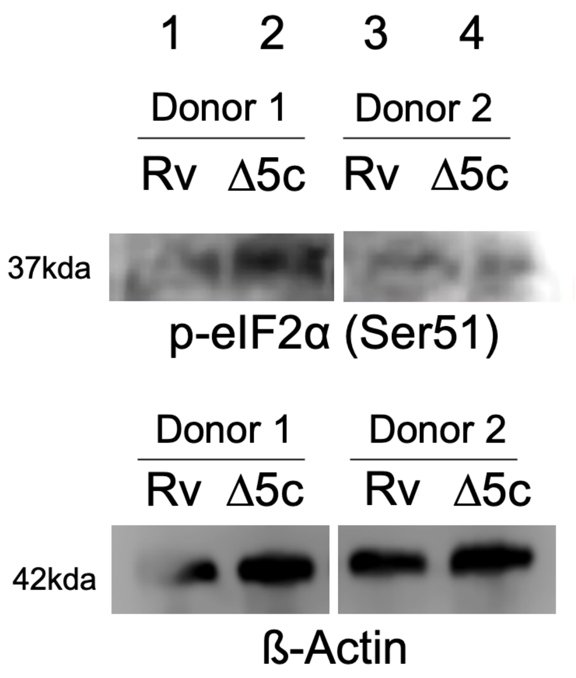
1 2 3 4
Donor 1
Donor 2
Rv
Δ5c
Rv
Δ5c
37kda
p-eIF2α (Ser51)
Donor 1
Donor 2
Rv
Δ5c
Rv
Δ5c
42kda
ß-Actin

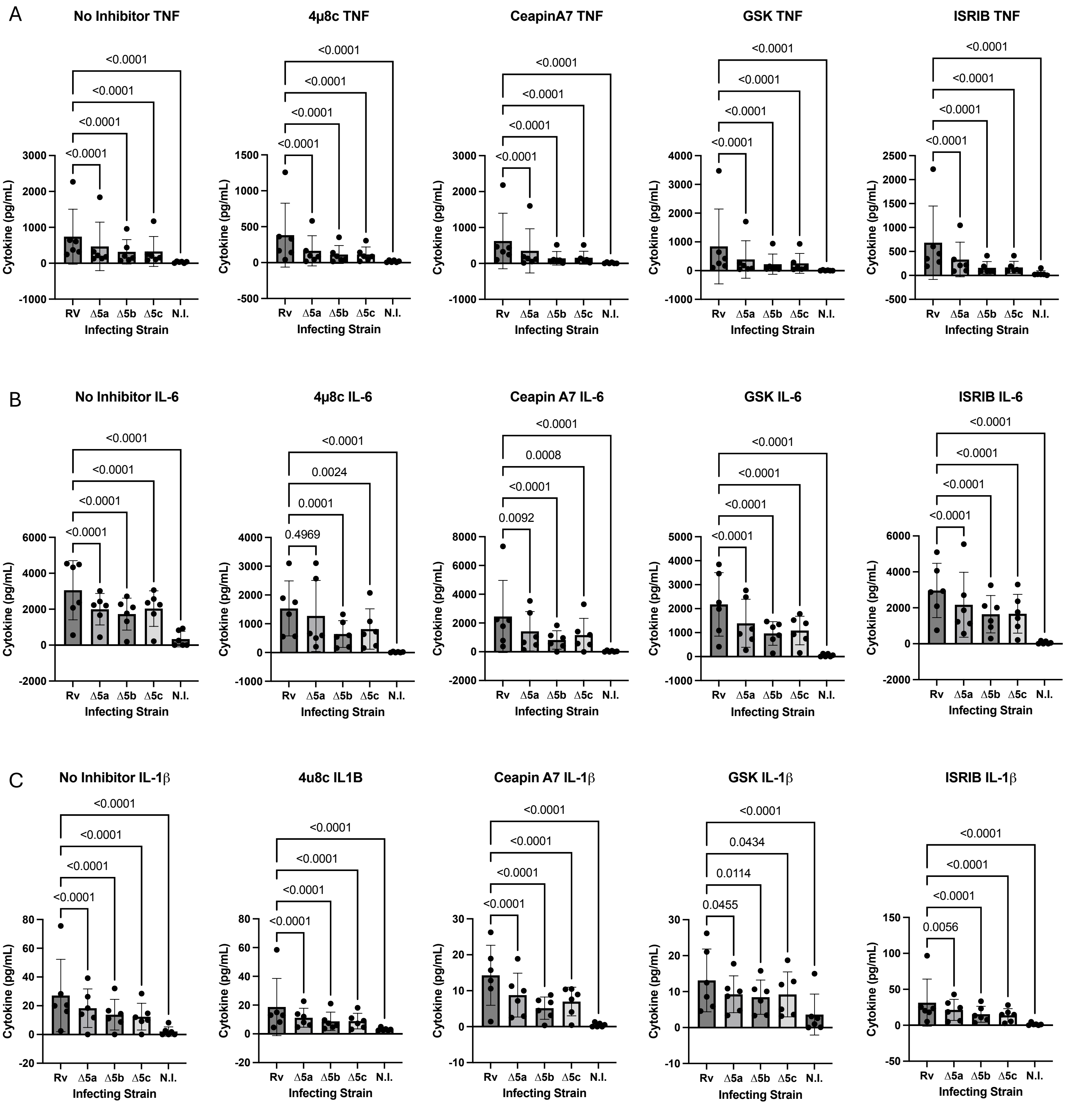

A
No Inhibitor TNF
<0.0001
<0.0001
<0.0001
<0.0001
4μ8c TNF
<0.0001
<0.0001
<0.0001
<0.0001
CeapinA7 TNF
<0.0001
<0.0001
<0.0001
<0.0001
GSK TNF
<0.0001
<0.0001
<0.0001
<0.0001
ISRIB TNF
<0.0001
<0.0001
<0.0001
<0.0001
B
No Inhibitor IL-6
<0.0001
<0.0001
<0.0001
<0.0001
4μ8c IL-6
<0.0001
0.0024
0.0001
0.4969
Ceapin A7 IL-6
<0.0001
0.0008
<0.0001
0.0092
GSK IL-6
<0.0001
<0.0001
<0.0001
<0.0001
ISRIB IL-6
<0.0001
<0.0001
<0.0001
<0.0001
C
No Inhibitor IL-1β
<0.0001
<0.0001
<0.0001
<0.0001
4u8c IL1B
<0.0001
<0.0001
<0.0001
<0.0001
Ceapin A7 IL-1β
<0.0001
<0.0001
<0.0001
<0.0001
GSK IL-1β
<0.0001
0.0434
0.0114
0.0455
ISRIB IL-1β
<0.0001
<0.0001
<0.0001
0.0056
Cytokine (pg/mL)
RV
Rv
Δ5a
Δ5b
Δ5c
N.I.
Infecting Strain

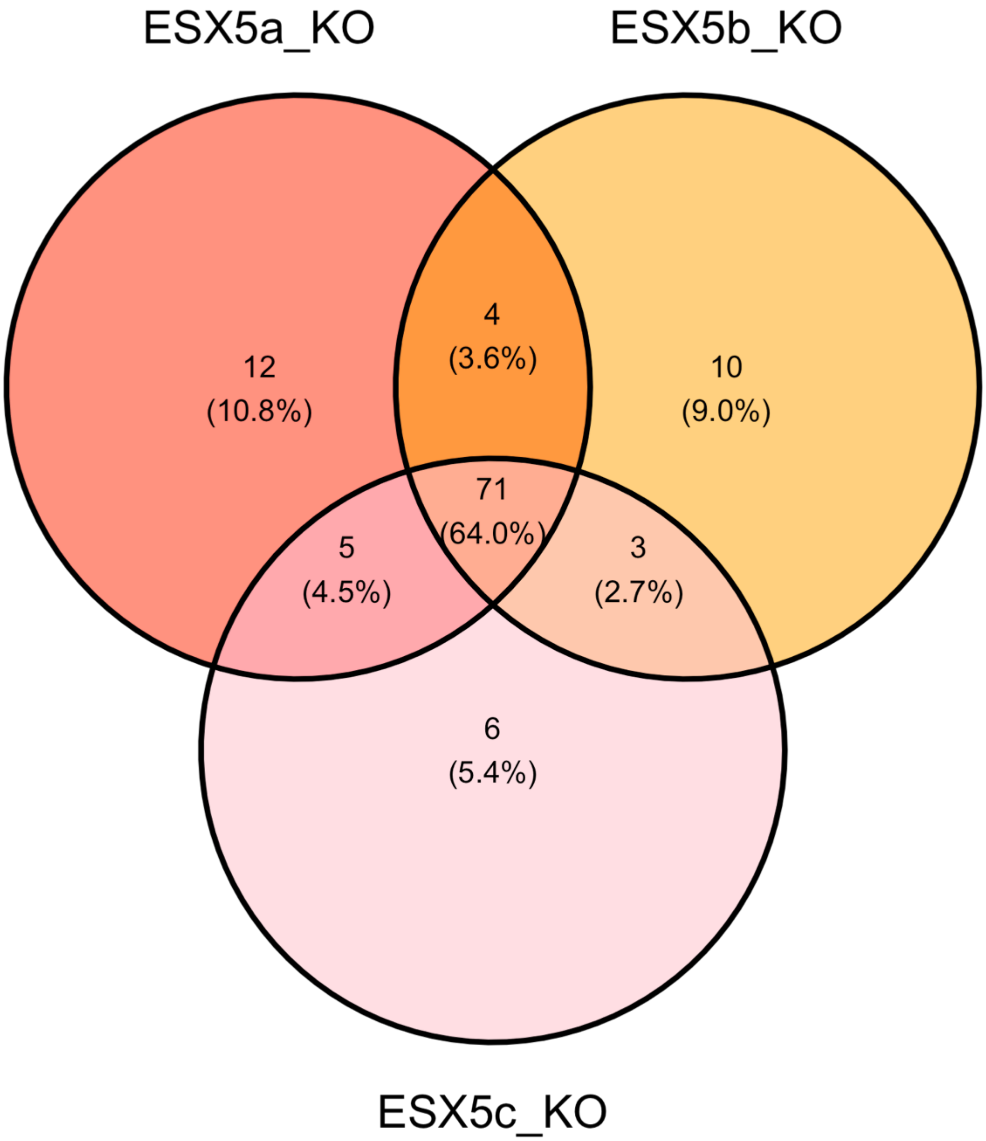
ESX5a_KO
ESX5b_KO
12
(10.8%)
4
(3.6%)
10
(9.0%)
71
(64.0%)
5
(4.5%)
3
(2.7%)
6
(5.4%)
ESX5c_KO

### Supplemental figures

**Supplemental Figure 1 – EccC5 CRISPRi knockdown efficiency and replication defect in macrophages.** *SF1A. EccC5 knockdown efficiency*. Bar chart depicts knockdown percentage (y-axis) of target gene for EccC5 (EccC5i +ATc) and the wildtype control (H37Rv +ATc) (x-axis). *SF1B. Broth culture EccC5i strain growth compared to wild type.* Bar chart shows the growth of EccC5i strain (y-axis, OD600) with and without inducer (x-axis, strain and ATc) for N=4 separate experiments. *SF1C. EccC5i strain is attenuated in human macrophages.* Bar charts display total CFU recovered (y-axis, log 10 CFU) from EccC5i induced (+ATc) and uninduced (-ATc) along with and H37Rv induced control (x-axis) across two donors (technical triplicate). *SF1B*. Statistics calculated using two-tailed T-test (95% CI, $p \leq 0.05$). *SF1C.* Statistics calculated using one way ANOVA with post-hoc T test using Dunnett correction (CI 95%, $p \leq 0.05$).

**Supplemental Figure 2 –ESX-5 paralog deletion validation.** *SF2A. PCR gel image for hygromycin resistance amplification.* Sixteen total isolates tested via PCR for hygromycin resistance gene (numbers indicate gel lane; red boxes indicate amplification fragment of ~2200 base pairs). *SF2B. qPCR of deleted gene targets.* Bar graph depicts fold change expression (y-axis) of ESX-5a, ESX-5b, and ESX-5c within respective deletion mutants (x-axis) run in N=3 separate experiments. Red dotted line indicates wildtype expression for respective gene target. *SF2C. Macrophage cell health is not ESX-5 dependent during in vitro infection.* Dot plots display the RFU measurement (y-axis) from cells treated with Alamar Blue and infected with varying strains of Mtb (x-axis, N.I. = uninfected, blnk = media only). Graphs depict respective time point measurements evaluating accumulated level of reduced Alamar Blue (saturation reached at 24hpi, decay observed at 48hpi). *SF2B-C.* Error bars show standard deviation of the mean.

**Supplemental Figure 3 – ESX-5 paralog complementation and macrophage cytokine phenotype**. *SF3A-C. ESX-5 complementation restores cytokine production compared to wild type macrophage infections.* Graphs display TNF, IL-6 and IL-1β levels across H37Rv, respective knockout strains, complement strains, and uninfected control. Bar charts show expression levels (y-axis, pg/mL) of respective cytokines across N=3 donors (technical triplicate) for a given comparison of WT, knockout, complement strain, and uninfected (N.I.) (x-axis). Statistics for all graphs calculated using two-way ANOVA with a post-hoc T-test and Dunnett correction for multiple comparisons (95% CI, $p \leq 0.05$). Error bars show standard deviation of the mean.

**Supplemental Figure 4 – ESX5 and macrophage 24-hour cytokine mRNA expression**. *SF4. Cytokine mRNA induction is not ESX-5 dependent at 24hrs post infection*. Bar charts represent the relative expression (y-axis, fold change expression) of TNF, IL-6, and IL-1β transcripts across N=3 donors (technical singlet) for respective infection conditions (x-axis). Statistics for all analyses generated using paired, two-tailed T-test (95% CI, $p \leq 0.05$). Error bars show standard deviation of the mean.

**Supplemental Figure 5 – Phospho- eIF2α (Ser51) protein levels in ESX-5 paralog mutant and wild type Mtb-infected cells.** *SF5. Western blot measurement of. phospho-eIF2α (Ser51) and control ß-Actin protein across lanes 1-4.* Lanes 1-2 shows protein from donor one, lanes 3-4 show donor 2. Lanes 1 and 3 show H37Rv conditions while lanes 2 and 4 show Δ5c condition.

**Supplemental Figure 6 – UPR and ISR inhibition does not rescue ESX-5 dependent cytokine phenotype.** *SF6A-C TNF, IL6, and IL-1β levels do not differ in an ESX5-dependent manner in macrophages treated with no inhibitor, 4μ8c (IRE1 inhibitor), Ceapin A7 (ATF6 inhibitor), GSK2606414 (PERK inhibitor), or ISRIB (p-eIF2 inhibitor) respectively*. *SF6A-C*. Graphs depict levels of given cytokine (y-axis, pg/mL) across N=6 donors (technical triplicate) under treatment with a respective inhibitor (Title) and bacterial strain (x-axis, N.I.= uninfected). Statistics for all analyses generated using two-way ANOVA with post-hoc T-test and Dunnett correction for multiple comparisons (95% CI, $p \leq 0.05$). Error bars show standard deviation of the mean.

**Supplemental Figure 7 – Overlap of Mtb ESX-5 paralog knockout strain transcriptional profiles**. SF7. Venn diagram shows three circles corresponding to significant gene expression for a given ESX-5 knockout compared to H37Rv wildtype. Gene distributions by circle and overlaps represent which genes are shared between which mutants and which are unique within respective strains. 71 (or 64%) of all genes identified are shared.

### Supplemental Tables

**Supplemental Table 1 – Mtb rRNA Depletion Probe Sequences.** Table shows custom probe sequences used for removing ribosomal RNA during Mtb library preparation.

**Supplemental Table 2- Macrophage miRNA profile from H37Rv and ∆ESX-5 paralog infections.** Table shows absolute counts for given strain and human miRNA combinations. Columns are organized by infecting strain (in biological triplicate). Rows are organized by miRNA species detected.

**Supplemental Table 3 – Differentially expressed genes of knockouts vs. wild type H37Rv in broth culture condition. Strains organized by sheets.** Tables show expression levels for all genes detectable in the knockouts as compared with H37Rv control across sheets. In columns, the table shows analysis outputs by gene. Column values "Gene", "log2FoldChange", and "padj" represent the gene name, log2 fold change in expression, and FDR corrected p-value, respectively. Significant genes for this data set were determined by setting a strict cutoff of Log2 Fold Change ≥1/≤-1 and an FDR corrected p-value ≥0.01.

**Supplemental Table 4 – Shared and unique genes across knockouts vs wildtype.** Table broken down by sheets showing genes that met significance cutoff across individual knockout strains, shared significant genes between all strains, and unique genes among individual knockouts (inclusion criteria: Log2 Fold Change ≤-1, FDR ≤0.01). All genes identified in column "Overlapping Genes" were shared across all strains and used for network analysis.
